# Mitochondrial genome-encoded mitomiRs regulate cellular plasticity and susceptibility to ferroptosis in triple-negative breast cancer

**DOI:** 10.1101/2024.11.16.623958

**Authors:** Amoolya Kandettu, Joydeep Ghosal, Jesline Shaji Tharayil, Raviprasad Kuthethur, Sandeep Mallya, Rekha Koravadi Narasimhamurthy, Kamalesh Dattaram Mumbrekar, Yashwanth Subbannayya, Naveena AN Kumar, Raghu Radhakrishnan, Shama Prasada Kabekkodu, Sanjiban Chakrabarty

## Abstract

Ferroptosis is a distinct form of regulated cell death promoted by iron-dependent lipid peroxidation. The metabolic plasticity of cancer cells determines their sensitivity to ferroptosis. Although mitochondrial dysfunction contributes to metabolic reprogramming in cancer cells, its role in ferroptosis remains to be identified. We identified that the mitochondrial genome encodes 13 miRNAs (mitomiRs) that are highly expressed in breast cancer cell lines and patient-derived tumor samples. Expression analysis revealed that mitomiRs are upregulated in basal-like triple-negative breast cancer (TNBC) cells compared to mesenchymal stem-like TNBC cells. Interestingly, 11 out of the 13 mitomiRs bind to the 3′UTR of zinc finger E-box-binding homeobox 1 (ZEB1), a transcription factor, involved in epithelial to mesenchymal transition (EMT) in breast cancer. Using mitomiR-3 mimic, inhibitor and sponges, we confirmed that mitomiR-3 indeed regulate ZEB1 expression in TNBC cells. Increased mesenchymal features in TNBC contributed to vulnerability to pro-ferroptotic metabolic reprogramming sensitizing to cell death in *in vitro* and *in vivo* models. Some of the challenges associated with pro-ferroptotic drugs includes lack of cancer cell specificity, low targeting ability, normal tissue toxicity contributing to their limited clinical application as cancer therapeutics. Here, we identified mitomiRs which are highly expressed in TNBC subtypes with low expression in normal breast cells making them an ideal candidate for selective inhibition for targeted therapy. Further, we demonstrated that the inhibition of mitomiRs in triple-negative breast cancer cells promote pro-ferroptotic metabolic reprogramming which can be exploited as novel vulnerability for targeted ferroptotic induction in cancer cells avoiding the normal tissue toxicity. Collectively, our results indicate a novel mechanism of mitochondrial miRNA mediated ferroptosis sensitivity in TNBC subtypes which could be exploited to develop potential miRNA-based therapeutics.

## INTRODUCTION

Triple-negative breast cancer (TNBC) is a heterogeneous and aggressive subtype of breast cancer, characterized by the absence of estrogen receptor (ER), progesterone receptor (PR), and human epidermal growth factor receptor 2 (HER2). Its molecular profile limits therapeutic options, because of the lack of targets for hormone therapies and HER2 inhibitors that are effective in other breast cancer subtypes. TNBC is categorized into six distinct molecular subtypes on the basis of gene expression status and cell type of origin, each with unique clinical outcomes and therapeutic responses ^1,2^. The basal-like subtypes (BL-1 and BL-2) are more epithelial in nature with characteristic features of high genomic instability and proliferative capacity^3^. Despite the relative chemosensitivity observed in some cases, BL-2 subtype tumors are more chemo refractory in nature ^4^. The mesenchymal stem-like (MSL) subtype is characterized by epithelial-mesenchymal transition (EMT), metabolic reprogramming, high invasiveness and resistance to conventional therapies ^3,5^. These subtypes present distinct cellular and metabolic phenotypes emphasizing the need for targeted, subtype-specific treatments. Multiple studies have reported the role of EMT in cancer invasion, migration and metastasis ^6^.

During EMT, cells can shift from an epithelial (E) state, which is characterized by the presence of cell-to-cell junctions, apical basal polarity, and interactions with the basement membrane, to a mesenchymal (M) state, which is characterized by a fibroblast-like appearance and enhanced migratory and invasive capabilities^7^. This process is largely regulated by a set of EMT-transcription factors (EMT-TFs) ZEB1, ZEB2, TWIST1, SNAI1, and SNAI2, whose expression is highly selective, and varies depending on the context and cell type ^8^. The zinc-finger E-box binding homeobox 1 (ZEB1) gene serves as a key regulator of EMT, and is considered a potential indicator of poor prognosis in TNBC patients ^9–11^.

Ferroptosis is a regulated form of cell death, characterized by the iron-dependent accumulation of lipid peroxides^12^. This process is triggered by the accumulation of iron and reactive oxygen species (ROS), leading to the oxidation of polyunsaturated fatty acids (PUFAs) within cell membranes, ultimately resulting in cell death. Unlike apoptosis, which is caspase-dependent, ferroptosis is driven by the disruption of cellular redox homeostasis due to inactivation of glutathione peroxidase 4 (GPX4), an enzyme crucial for detoxifying lipid peroxides^13^. Undifferentiated cells and cells with a partial mesenchymal phenotype are highly susceptible to ferroptosis and are activated by the EMT-TF ZEB1 which induces the expression of PUFA biosynthesis enzymes and concurrently inhibits monounsaturated fatty acids (MUFAs) synthesis^14^. PUFAs are considered the main facilitators of ferroptosis because of their enrichment in cell membranes and vulnerability to lipid peroxidation^15,16^. The enrichment of PUFAs in the cell membrane promotes sensitivity to ferroptosis^17,18^. Previous studies have indicated that mesenchymal-type cancer cells have increased PUFA metabolism which increases their vulnerability to ferroptosis^19^. Mitochondria play a central role in lipid and redox metabolism. Earlier studies have revealed the role of mitochondrial electron transport activity in ROS generation and ferroptosis has been demonstrated ^20^. The identification of the cellular and metabolic changes that promote ferroptosis sensitivity in triple-negative breast cancer holds great promise as a novel vulnerability that can be targeted for therapeutic purposes.

Mitochondrial genome encodes 13 microRNAs (mitomiRs), that are differentially expressed in breast cancer cell lines and tissue specimens^21^. Interestingly, almost all mitomiRs are highly expressed in basal-like TNBC (MDA-MB-468) and significantly downregulated in mesenchymal-like TNBC (MDA-MB-231)^21^. In the present study, we showed that 11 out of the 13 mitomiRs target the EMT-TF ZEB1 gene, indicating their possible role in the regulation of breast cancer cell plasticity and EMT. Through biochemical and genomic analyses, we observed that mitomiR-3 indeed regulates ZEB1 expression in TNBC cells. By sponging mitomiR-3 in TNBC cells, we observed that the loss of mitomiR-3 promoted ZEB1 mediated transcriptional repression of GPX4, a key enzyme, that facilitates ferroptosis protection by converting lipid hydroperoxides to lipid alcohols. MitomiR-3 sponge cells showed increased ferroptotic cell death in TNBC cell line and *in vivo* mouse tumors. Gene expression and metabolomics analysis confirmed that the inhibition mitomiR-3 in TNBC cells promote pro-ferroptotic gene expression and increased the levels of PUFA metabolites, which contributed to lipid peroxidation and ferroptosis. Collectively, these evidence provide opportunity for selective targeting of mitomiRs in cancer cells which are vulnerable to ferroptotic cell death without damaging the normal cells in the complex tumor microenvironment.

## RESULTS

### mitomiR-3 regulates ZEB1 expression in TNBC cell lines

Mitochondria plays a central role in metabolic reprogramming in breast cancer. Previously, we have demonstrated that the mitochondrial genome encodes 13 miRNAs that are differentially expressed in breast cancer cells^21^. We have shown that mitomiRs translocate to the cytoplasm and are involved in posttranscriptional regulation of nuclear genome-encoded genes by binding to the RISC complex protein Ago2 in the cytoplasm^21^. Expression analysis of selected mitomiRs revealed that mitomiRs are upregulated in basal-like TNBC cells (MDA-MB-468) compared to mesenchymal stem-like TNBC cells (MDA-MB-231). Interestingly, bioinformatic analysis for target prediction identified that 11 out of the 13 mitomiRs bind to 3′-untranslated region (3′-UTR) of the Zinc-finger E-box binding homeobox 1 (ZEB1) gene, a known epithelial-mesenchymal transition (EMT) associated transcription factor (Supplementary Figure 1). One of the mitomiRs, mitomiR-3, which targets ZEB1, interacts with the RISC complex protein Ago2 inside mitochondria and in the cytoplasm suggesting its potential role in the regulation of gene expression (Fig. 1a, b). MitomiR-3, a mitochondrial genome-encoded miRNA, is highly expressed in MDA-MB-468 when compared to MCF-7, MDA-MB-231 and MCF10A (Fig. 1c). Expression analysis of mitomiR-3 in normal and breast tumor tissues showed increased mitomiR-3 expression in the tumor tissues of breast cancer patient samples (Figure 1d, Supplementary Table 1). Bioinformatic analysis revealed that mitomiR-3 seed sequence binds to miRNA response element (MRE) on the ZEB1 3′-UTR (Fig. 1e). The seed sequence of mitomiR-3 was conserved across species suggesting its possible regulatory role in non-human model organisms (Fig. 1e).

**Fig. 1.**
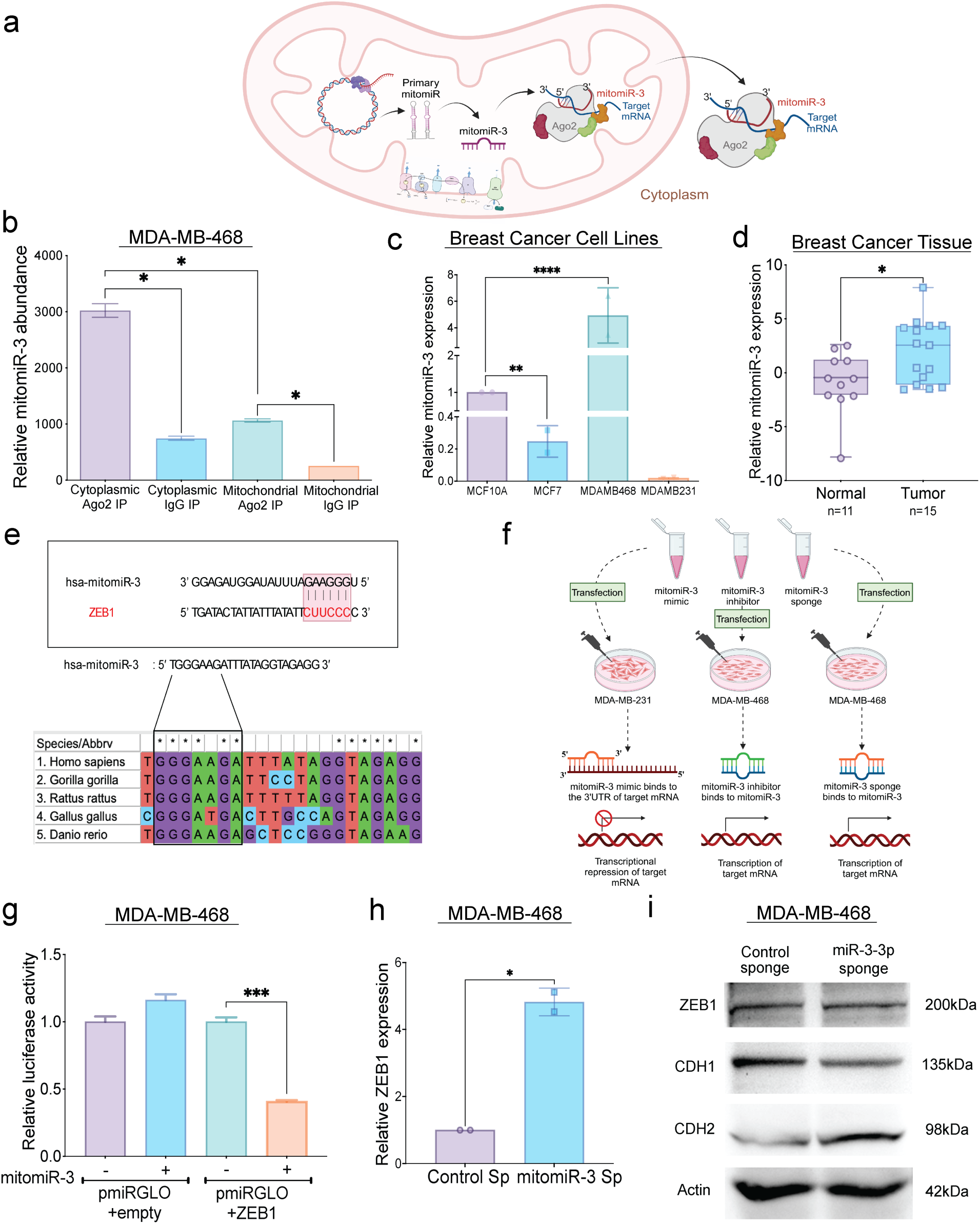
Mitochondrial genome-encoded microRNA-3 (mitomiR-3) regulates ZEB1 expression in TNBC cells. a) Schematic representation of mitomiR-3 biogenesis and its association with Ago2 in the cytoplasm and within mitochondria. mitomiR-3 is encoded from the 16S rRNA region on the mitochondrial DNA. b) Relative mitomiR-3 abundance in Ago2 IP elute of cytoplasmic and mitochondrial fractions. IgG rabbit Ab was used as experimental control and empty beads were used as negative control. Fold change was calculated by normalizing the Ago2-IP and IgG-IP elute amplification to empty beads amplification. mitomiR-3 enrichment was observed in both mitochondrial and cytoplasmic Ago2 elute, indicating mitomiR-3 association with mitochondrial RISC fraction and cytoplasmic RISC fraction. c) Relative mitomiR-3 expression in MCF-7, MDA-MB-468 & MDA-MB-231 cell lines, in comparison to MCF10A cell line. 5S rRNA was used as endogenous control and relative quantity was calculated using the formula 2^-ΔΔCт^. MDA-MB-468 (Basal-like TNBC) shows increased expression of mitomiR-3, while MDA-MB-231 (Mesenchymal stem-like TNBC) shows significantly reduced mitomiR-3 expression. d) Relative mitomiR-3 expression in normal and breast cancer tissue specimen. RNU6B was used as the internal control and relative quantity was calculated using the formula 2^-ΔCт^. Breast cancer tumor specimen showed increased mitomiR-3 enrichment in comparison to normal breast tissue specimen. e) *In-silico* analysis predicted nuclear gene *ZEB1* as the putative target of mitomiR-3. The seed region of mitomiR-3 binds to the miRNA response element (MRE) region on the *ZEB1* 3’-UTR. mitomiR-3 seed region is conserved across vertebrates. f) Schematic workflow of mitomiR-3 overexpression in TNBC cells using mimic, and inhibition of mitomiR-3 by using inhibitor and sponge transfection and mitomiR-mRNA targeting. g) Dual-Luciferase reporter assay upon 48-hour co-transfection with 3′UTR of *ZEB1* mRNA sequence cloned pmirGLO vector along with mitomiR-3 mimic or negative control oligos in MDA-MB-468 cell line. Relative luciferase activity was calculated upon normalizing the activity of luciferase enzyme to *Renilla*-luciferase enzyme activity. (+) and (−) signs indicate the presence and absence of mitomiR-3 mimic respectively. The luciferase activity was reduced in pmiRGLO+*ZEB1* construct transfected with mimic while there was no change observed in the pmiRGLO empty vector construct. h) qRT-PCR analysis showing relative *ZEB1* mRNA expression in mitomiR-3 Sp cells compared to Control Sp cells. Expression of *ZEB1* in mitomiR-3 Sp was significantly upregulated when compared to Control Sp. i) Western blot analysis of ZEB1, CDH1 and CDH2 protein expression in Control Sp and mitomiR-3 Sp cell lines. mitomir-3 Sp cells showed increased protein expression of ZEB1 and CDH2 (mesenchymal marker), while protein expression of CDH1 (epithelial marker) was reduced. \**p-value* <0.05, ** *p-value* <0.01, *** *p-value* <0.001 and **** *p-value* <0.0001.

To experimentally validate the post-transcriptional regulation of ZEB1 by mitomiR-3, we employed multiple strategies including the transfection of a chemically modified mitomiR-3 mimic in MDA-MB-231 cells (mesenchymal stem-like TNBC cells with low mitomiR-3 and high ZEB1 expression) (Fig. 1f). Furthermore, we transfected a mitomiR-3 inhibitor into MDA-MB-468 (basal-like TNBC cells with high mitomiR-3 and low ZEB1 expression) cells which inhibits the mitomiR-3 function via competitive interactions with the mitomiR-3 seed region, thereby preventing the interactions with the target gene, ZEB1 (Fig. 1f). To create a stable mitomiR-3 loss-of-function sponge model, a retroviral-based mitomiR-3 sponge expressing MDA-MB-468 cell line was generated. MitomiR-3 sponge cells produce a mitomiR-3 antisense sequence that competitively binds to endogenous mitomiR-3 and prevents mitomiR-3/target gene interactions (Fig. 1f). The binding of mitomiR-3 to the ZEB1 3′-UTR was confirmed by dual-luciferase reporter assay which revealed that, compared with the control, the mitomiR-3 mimic repressed ZEB1 reporter activity (Fig. 1g). Inhibition of mitomiR-3 by sponge promoted ZEB1 upregulation in TNBC cells confirming that mitomiR-3 negatively regulate ZEB1 expression in TNBC cells (Figure 1h). ZEB1 is a transcriptional repressor of the epithelial cell marker gene CDH1 (E-cadherin) in cancer cells^10,22^. We observed that ZEB1 upregulation in mitomiR-3 sponge cells led to decreased CDH1 expression and increased expression of the mesenchymal marker gene CDH2 (N-cadherin) (Fig. 1i).

### mitomiR-3 sponge cells showed mesenchymal morphology

Immunofluorescence analysis confirmed increased ZEB1 nuclear localization in mitomiR-3 sponge cells compared to control cells (Fig. 2a, b). ZEB1 expression in TNBC cells promotes epithelial-to-mesenchymal transition (EMT) and stemness^8^. We asked whether mitomiR-3 expression contributes to the maintenance of the epithelial cell phenotype in TNBC cells (Fig. 2c). Gene expression analysis using RNA sequencing data from negative control, mitomiR-3 mimic (upregulation) transfected MDA-MB-231 and mitomiR-3 sponge (loss-of-function) transfected MDA-MB-468 cells identified increased expression of ZEB1 and other mitomiR-3 target genes involved in EMT pathway in mitomiR-3 sponge cells (Fig. 2d, Supplementary Table 2). EMT pathway genes upregulated in mitomiR-3 sponge cells were further analyzed in publicly available breast cancer cell line RNAseq datasets from DSMZ CellDive (Supplementary Fig. 2). Interestingly, EMT pathway genes upregulated in mitomiR-3 sponge cells (Fig. 2d) were highly expressed in TNBC cell lines with mesenchymal features (MDA-MB-231, Hs-578T, CAL-120) (Supplementary Fig. 2). Gene ontology and pathway analysis showed enrichment of the TGF-β signaling pathway, and the EMT pathway in mitomiR-3 sponge cells (Supplementary Fig. 3). Next, to assess whether mitomiR-3 expression alters the cellular phenotype of TNBC cells, imaging analysis was performed using mitomiR-3 mimic transfected MDA-MB-231 cells, mitomiR-3 inhibitor and mitomiR-3 sponge transfected MDA-MB-468 cells. We observed that mitomiR-3 mimic transfected cells (MDA-MB-231) presented increased epithelial cell morphology (Supplementary Fig. 4a). Similarly, the inhibition of mitomiR-3 in mitomiR-3 inhibitor transfected cells (MDA-MB-468) resulted in increased mesenchymal cell morphology (Supplementary Fig. 4b). ZEB1 mediated EMT promotes actin cytoskeleton remodeling in cancer cells^23^. We observed that, compared with negative control cells, phalloidin-stained mitomiR-3 mimic transfected cells (MDA-MB-231) presented decreased filopodial extensions suggesting increased epithelial features (Fig. 2e-g, i). Similarly, compared with negative control cells, mitomiR-3 inhibitor transfected, and sponge cells (MDA-MB-468) presented increased mesenchymal features (Fig. 2e, h and Supplementary Fig. 4c).

**Fig. 2.**
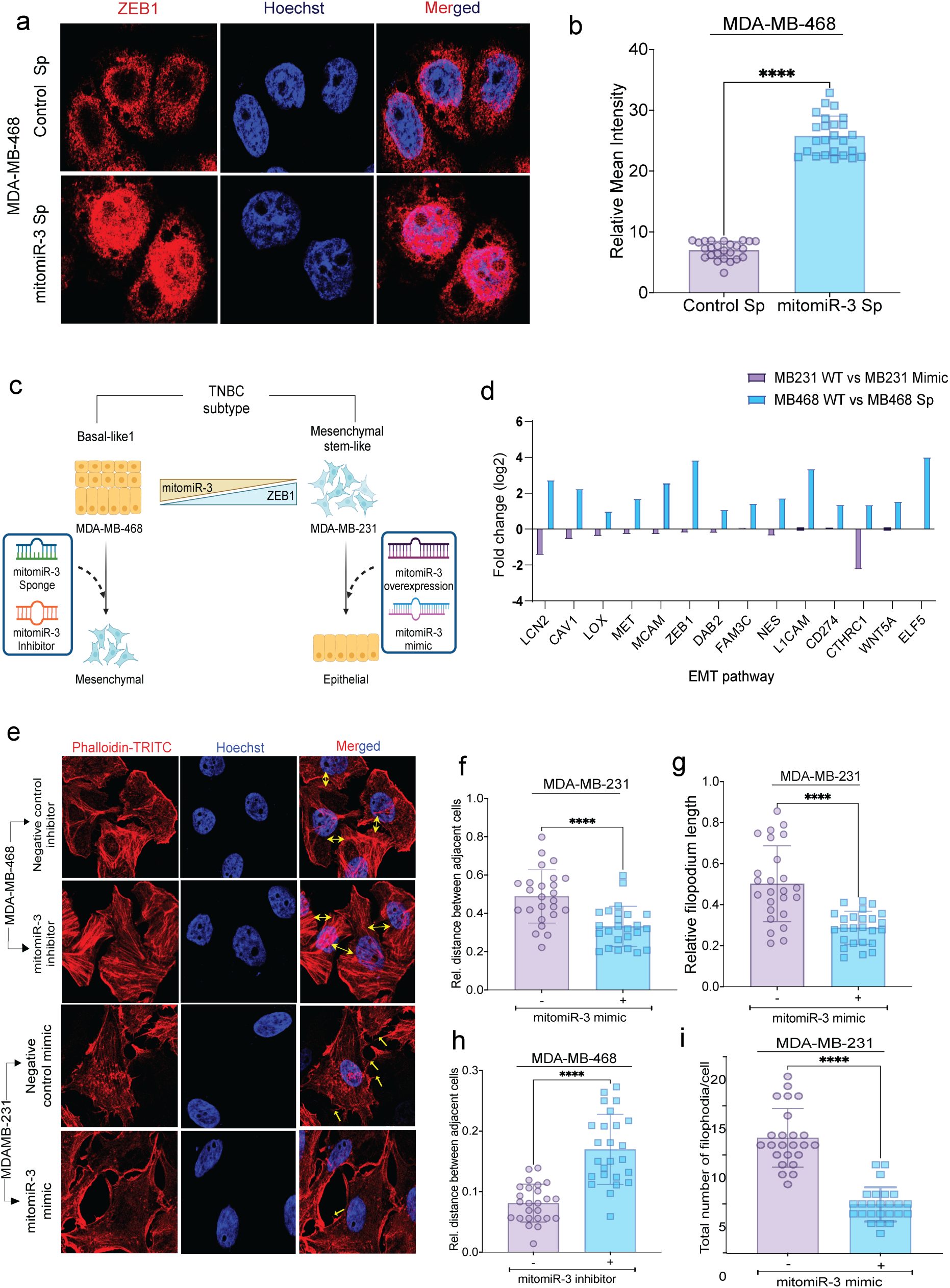
Inhibition of mitomiR-3 promote mesenchymal phenotype in MDAMB-468 cells. a) Immunofluorescence imaging of ZEB1 in Control Sp and mitomiR-3 Sp cells co-stained with Hoechst. Images were acquired using 64x oil immersion objective of confocal microscope (Scale bars, 50μm). A total of 15 different ROIs were analyzed per group. b) Bar graph showing the relative mean intensity of ZEB1 in Control Sp and mitomiR-3 Sp cells. mitomiR-3 Sp cells showed increased nuclear localization of ZEB1, while the control Sp cells showed no visible nuclear localization. c) Schematics showing morphological variation between TNBC cell lines, MDA-MB-468 (BL-1) and MDA-MB-231 (MSL). The schematics hypothesizes, blocking the mitomiR function (mitomiR-3 sponge/ mitomiR-3 inhibitor) can shift the cellular phenotype of MDA-MB-468 (BL-1) to mesenchymal phenotype, while promoting (mitomiR-3 mimic/overexpression) its function can shift cellular phenotype of MDA-MB-231 (MSL) to epithelial phenotype. d) Bar graph depicting differential gene expression of key mesenchymal markers that showed upregulated expression in mitomiR-3 sponge transfected MDA-MB-468 cells and downregulated expression in mitomiR-3 mimic transfected MDA-MB-231 cells. e) Representative actin phalloidin imaging of TNBC cell lines, MDA-MB-468 and MDA-MB-231 transfected with 50nM of mitomiR-3 inhibitor, mimic and its respective negative control oligonucleotides (Scale bars, 50μm). Total of 14 different ROIs were analysed per group. f-i) Bar graph showing the relative distance between adjacent cells, average length and number of filopodia. Increased epithelial features were observed in mitomiR-3 mimic transfected cells which showed reduced filopodial length, number and distance between adjacent cells, while increased mesenchymal features were observed in mitomiR-3 inhibitor transfected cells. *\*p*<0.05, *\*\*p*<0.01, *\*\*\*p*<0.001 and *\*\*\*\*p*<0.0001.

### mitomiR-3 sponge inhibits cell proliferation, migration, invasion and promotes cell death in TNBC

High ZEB1 expression and EMT are associated with increased cell invasion, migration and metastasis. Interestingly, we observed that the inhibition of mitomiR-3 by sponge perturbs cell proliferation in TNBC cells (Fig. 3a). Analysis of S-phase cells by EdU staining in negative control and mitomiR-3 sponge cells confirmed significant decrease in S-phase cells in mitomiR-3 sponge cells than negative control cells (Fig. 3b-c). Furthermore, analysis of apoptosis by annexin V-FITC/PI staining showed increased cell death in mitomiR-3 sponge cells compared to negative control cells confirming that inhibition of mitomiR-3 function contribute to cell death in TNBC cells (Fig. 3d-g, Supplementary Fig. 5). ZEB1 upregulation in TNBC cells promotes increased invasiveness, cell migratory phenotype and metastasis. However, cell-based assays showed significant reduction in cell invasion and migratory phenotype in mitomiR-3 sponge cells (Fig. 3h-l). Further, analysis of *in vivo* tumor growth in control and mitomiR-3 sponge MDA-MB-468 cell line derived tumor xenograft (CDX) mouse models revealed a significant reduction in tumor volume in mitomiR-3 sponge cells derived CDX (Fig. 3m-o). Histology analysis by H&E staining in control and mitomiR-3 sponge mice confirmed increased cell death in mitomiR-3 sponge cells (Fig. 3p). Immunohistochemistry analysis confirmed increased ZEB1 expression and decreased cell proliferation marker Ki67 expression in mitomiR-3 sponge cells (Fig. 3p). Prussian blue staining is commonly used to study the intracellular iron content and ferroptosis in tissue specimens. Histopathology analysis showed significant increase in the positive area of Prussian blue staining in mitomiR-3 sponge derived CDX mouse tumors when compared to negative control cell derived CDX mouse tumors (Fig. 3p-q). Further, TUNEL staining showed altered tumor cell morphology and increased TUNEL positive tumor cells confirming ferroptotic cell death in mitomiR-3 sponge derived CDX mouse tumors when compared to control cell derived CDX mouse tumors (Fig. 3r-s).

**Fig. 3.**
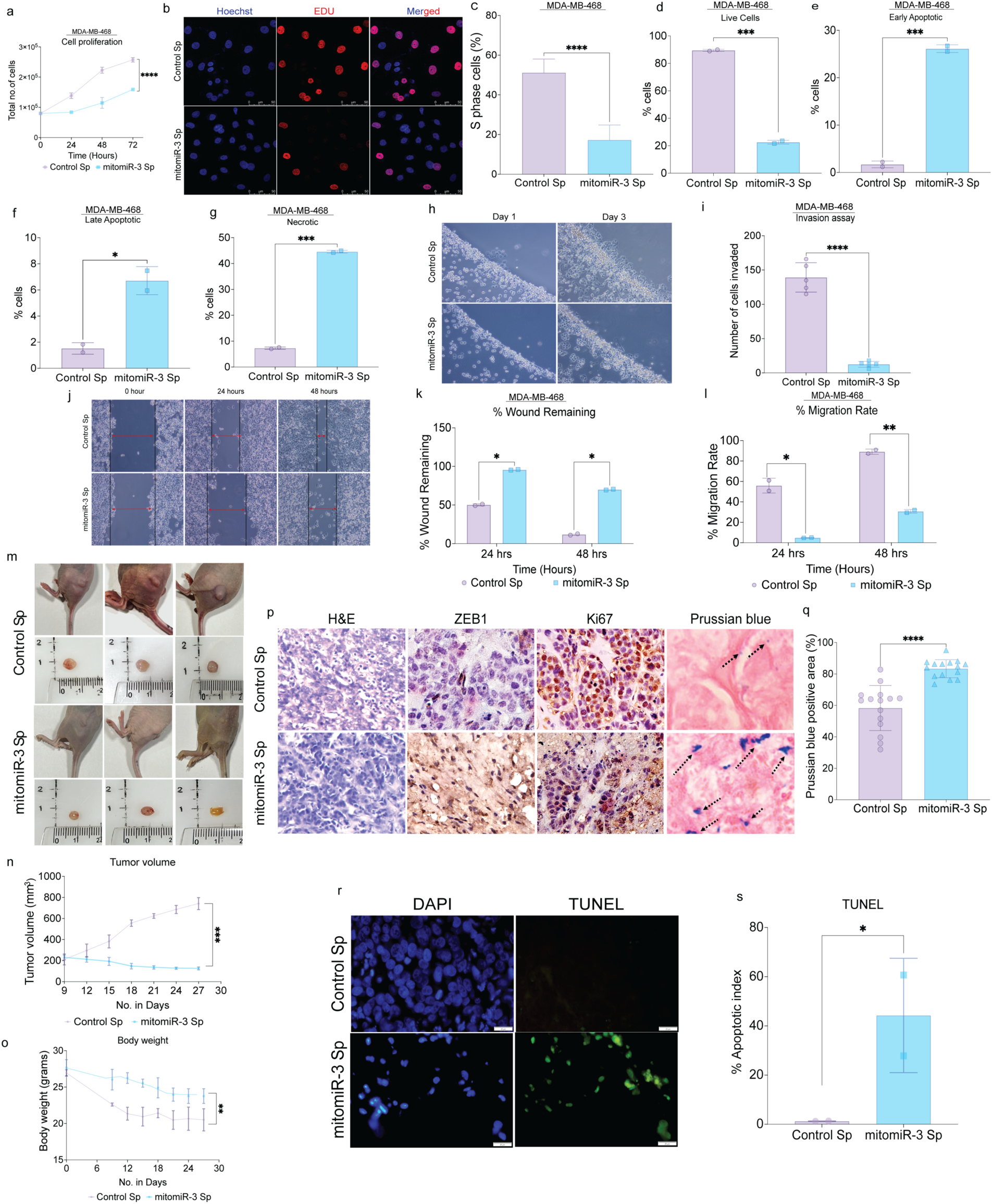
Inhibition of mitomiR-3 function perturbs cell proliferation, tumor growth and cell death and induces ferroptosis in MDA-MB-468 cells. a) Line graph showing the proliferation rate of mitomiR-3 Sp cells in comparison with Control Sp cells. mitomiR-3 Sp cells show a decrease rate of cell proliferation. b, c) Cell proliferation showed by EdU staining. S phase cells detected via EdU (red) and Hoechst-stained nuclei with blue (Scale bars, 5μm). A total of 15 different ROIs were analyzed per group. The percentage of cells in the S-phase were significantly reduced in mitomiR-3 Sp cells. d-g) Flow cytometry analysis for apoptosis analysis in Control Sp and mitomiR-3 Sp transfected MDA-MB-468 cells. The total percentage of live cells in control Sp group was ∼80% and mitomiR-3 Sp group was ∼80%. The total percentage of early and late apoptotic cells in control Sp group was ∼3% and ∼1.5% and mitomiR-3 Sp group was ∼35% and ∼7%. The total percentage of necrotic cells in control Sp group was ∼9% and mitomiR-3 Sp group was ∼45%. h) Bright field microscopic images of cell invasion by agarose spot assay (×20 magnification). i) Bar graph showing the number of cells invading the agarose spot. mitomiR-3 Sp cells showed reduction in their invasive capacity. j) Bright field microscopic images of wound healing assay (×20 magnification). k) Bar graph showing the percentage wound remaining. l) Bar graph showing the percentage migration rate. mitomiR-3 Sp cells show decreased rate of cell migration. m) Images of tumor growth in nude mice injected subcutaneously with Control Sp and mitomiR-3 Sp cell line derived xenografts (CDX). n) Tumor growth curve of Control Sp and mitomiR-3 Sp expressing MDA-MB-468 cells shows decrease in tumor volume in mitomiR-3 Sp CDX. o) Line graph showing the change in body weight in Control Sp and mitomiR-3 Sp CDX. p) Representative images of H&E, ZEB1 (Strong nuclear expression of ZEB1 protein seen in mitomiR-3 Sp CDX compared to control Sp CDX) and Ki67 (Brown nuclear stain highlighting Ki67 positive tumor cells was significantly less in mitomiR-3 Sp CDX compared to control Sp CDX) immunohistochemistry (×20 magnification) and Prussian blue staining (Prussian blue positive cells depict higher concentration of iron deposits in mitomiR-3 Sp CDX. Blue: iron deposits, Pink: cytoplasm) showing iron content in cell line-derived xenografts (CDX). q) Bar graph for quantification of Prussian blue positives cells. The Prussian blue positive intensity was calculated by the formula integrated optical density (IOD)/area. r) Representative image of TUNEL staining in mouse tissue samples. A TUNEL assay revealed apoptotic-positive cells marked by green staining and intact DNA marked by blue DAPI stain. s) Bar graph estimating the percentage apoptotic index showing mitomiR-3 Sp CDX have higher cell death compared to the control CDX. Total of 1000 cells were counted for each specimen from three different ROI. The apoptotic index was calculated by the formula apoptotic index (%) = 100 x TUNEL positive cells/ total cells. Data represented as mean ±SEM, *p<0.05. Scale bar, 100 µm for H&E images and 20µm for DAPI and TUNEL images. *p<0.05, **p<0.01, ***p<0.001 and ****p<0.0001.

### mitomiR-3 sponge induce pro-ferroptotic metabolic pathway gene expression in TNBC

We identified that mitomiR-3 targets multiple genes related to pro-ferroptotic lipid peroxidation, iron metabolism, oxidative stress and EMT pathways (Fig. 4a). To confirm the mitomiR-3 mediated regulation of pro-ferroptotic gene expression in TNBC, we performed RNA sequencing and metabolomics analysis using negative control, mitomiR-3 mimic and mitomiR-3 sponge transfected TNBC cell lines (MDA-MB-231 and MDA-MB-468) (Fig. 4b). Gene expression analysis of mitomiR-3 target genes in negative control (MDA-MB-231) and mitomiR-3 mimic transfected (MDA-MB-231) with negative control (MDA-MB-468) and mitomiR-3 sponge transfected (MDA-MB-468) TNBC cells showed increased expression of pro-ferroptotic genes involved in iron metabolism (LCN2, MYCN, HMGB1), EMT (ZEB1, TGFB), oxidative stress (KEAP1, CAV1, DNAJB6, SLC38A1, and MTDH1), and lipid metabolism (LOX, ALOX15, ALOX15B, ACSL1, and EGLN2) upon inhibition of mitomiR-3 in mitomiR-3 sponge cells (Fig. 4c). Pathway analysis of mitomiR-3 target genes showed increased expression of arachidonic acid metabolism genes (ALOX15, ALOX15B), alpha-linolenic acid metabolism genes (PLA2G4B, PLB1), linoleic acid metabolism (PLA2G4B, PLB1, ALOX15) and unsaturated fatty acid metabolism (FADS1, CYP4F3, PLA2G4B, ALOX15, and ALOX15B) in mitomiR-3 sponge cells (Fig. 4d). Arachidonic acid 15-lipoxygenase-1 (ALOX15) is an enzyme that promotes peroxidation of PUFA-phospholipids to generate bioactive metabolites to increase ferroptotic cell death.^24,25^ Recent studies have shown that mesenchymal cell types promote reprogramming of lipid metabolism and the polyunsaturated fatty acid (PUFA) biosynthesis pathway^14^. Inhibition of mitomiR-3 showed increased mesenchymal features in TNBC cells. Therefore, we wanted to know whether mitomiR-3 sponging will promote PUFA metabolism in TNBC cells. Indeed, metabolomics analysis of negative control and mitomiR-3 sponge TNBC cells showed increased PUFA metabolites arachidonic acid (AA), Dihomo-gamma-linolenic acid (DGLA), Linoleic acid (LA) and Eicosapentaenoic acid (EPA) in mitomiR-3 sponge cells (Fig. 4e). Pathway analysis of metabolites identified significant enrichment of alpha-linolenic acid metabolism, unsaturated fatty acid metabolism, sphingolipid metabolism, and linoleic acid metabolism in mitomiR-3 sponge cells confirming the role of mitomiR-3 loss-of-function in reprogramming of pro-ferroptotic metabolism in TNBC cells (Supplementary Fig. 6). The enrichment of PUFA metabolites in cancer cell promotes lipid peroxidation and ferroptosis^16,18^. Membrane phospholipids containing arachidonic acid is more prone to peroxidation promoting ferroptosis^26^. Previous studies have shown that redox-active iron forms highly reactive hydroxyl radicals via Fenton reaction to initiate lipid peroxidation^12,27–29^. Gene expression analysis of mitomiR-3 mimic and sponge cells confirmed the increased expression of iron metabolism genes in mitomiR-3 sponge cells (Fig. 4c, Supplementary Fig. 7a). Analysis of intracellular iron content using inductively coupled plasma-mass spectrometry (ICP-MS) in control and mitomiR-3 sponge cells showed significant increase in intracellular iron content in mitomiR-3 sponge cells (Supplementary Fig. 7b). Taken together, these data conclusively suggest that mitomiR-3 expression is critical for the regulation of iron and lipid metabolism in TNBC cells to evade ferroptotic cell death.

**Fig. 4.**
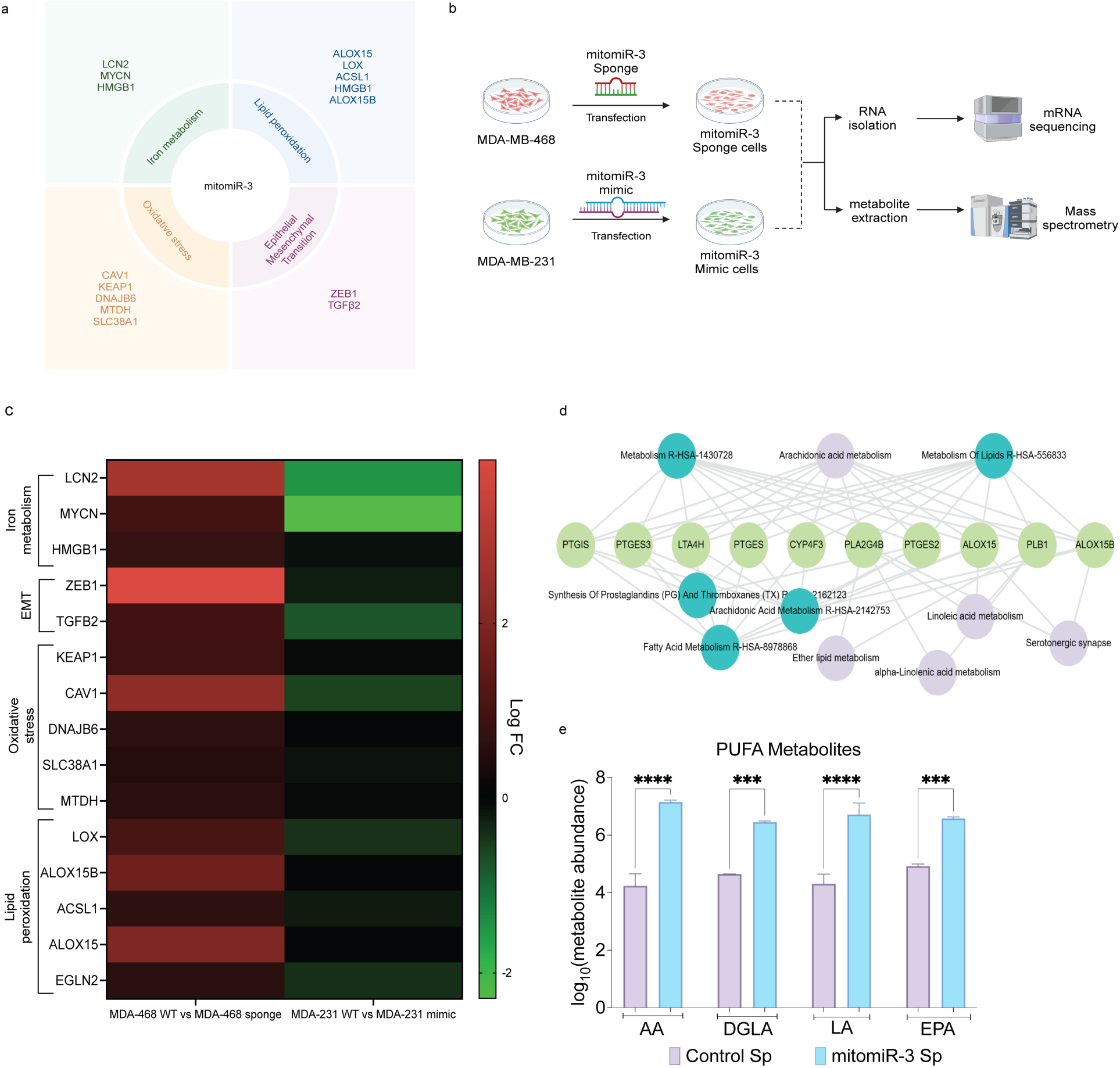
Inhibition of mitomiR-3 by sponge promotes upregulation of pro-ferroptotic gene expression and increase abundance of pro-ferroptotic metabolites in TNBC cells. a) Graphical representation of mitomiR-3 target genes functioning as ferroptosis driver genes b) Schematics of experimental workflow for RNA sequencing and mass spectrometric analysis for mitomiR-3 sponge and mimic transfected cells.. c) Heatmap of differential gene expression (log Fold Change) of ferroptosis driver genes that are predicted targets of mitomiR-3 in mitomiR-3 mimic transfected and sponge cells. d) Enriched pathway analysis of upregulated genes in mitomiR-3 LOF cells showed network of enriched PUFA metabolism genes. e) Fold change in metabolite abundance associated with PUFA in mitomiR-3 Sp and Control Sp cells based on mass spectrometric analysis. AA, Arachidonic acid; DGLA, Dihomo-gamma-linolenic acid; LA, Linoleic acid; EPA, Eicosapentaenoic acid. *p<0.05, **p<0.01, ***p<0.001 and ****p<0.0001.

### ZEB1 upregulation in mitomiR-3 sponge cells inhibits GPX4 expression in TNBC cells

Glutathione peroxidase 4 (GPX4) plays a central role in the cellular ferroptosis defense system. Depletion of GPX4 perturbs the reduction of phospholipid peroxides and promote ferroptotic cell death^13,24^. ZEB1 is a master transcription regulator and binds to the E-box motif present in more than 2000 gene promoters to either promote or inhibit specific gene expression^10^. We identified ZEB1 E-box binding motif (CACCTG) in the GPX4 promoter sequence (Fig. 5a). ZEB1 binding to GPX4 promoter was confirmed by chromatin immunoprecipitation (ChIP) assay using anti-ZEB1 antibody in combination with primers amplifying GPX4 promoter E-box sequence (Figure 5b-c). Further, ZEB1 upregulation contributed to transcriptional repression of GPX4 in mitomiR-3 sponge cells (Fig. 5d). This observation was further confirmed by GPX4 immunofluorescence analysis (Fig. 5e-f) and immunoblot analysis in control and mitomiR-3 sponge cells (Fig. 5g). ZEB1 is highly expressed in mesenchymal type breast cancer cell lines (Fig. 5h). GPX4 inhibition in mesenchymal type breast cancer cells promote lipid peroxidation^14^. High ZEB1 expression negatively correlated with GPX4 expression in breast cancer cell lines from CCLE database^30^. Further, analysis of public datasets for metastatic breast cancer^31^, we observed that majority of breast cancer patient with high ZEB1 expression showed low GPX4 gene expression indicating regulatory role of ZEB1 in GPX4 inhibition promoting ferroptosis sensitivity (Fig. 5j). These findings suggest that mitomiR-3 sponging inhibit GPX4 expression via ZEB1 upregulation contributing to ferroptosis in TNBC cells.

**Fig. 5.**
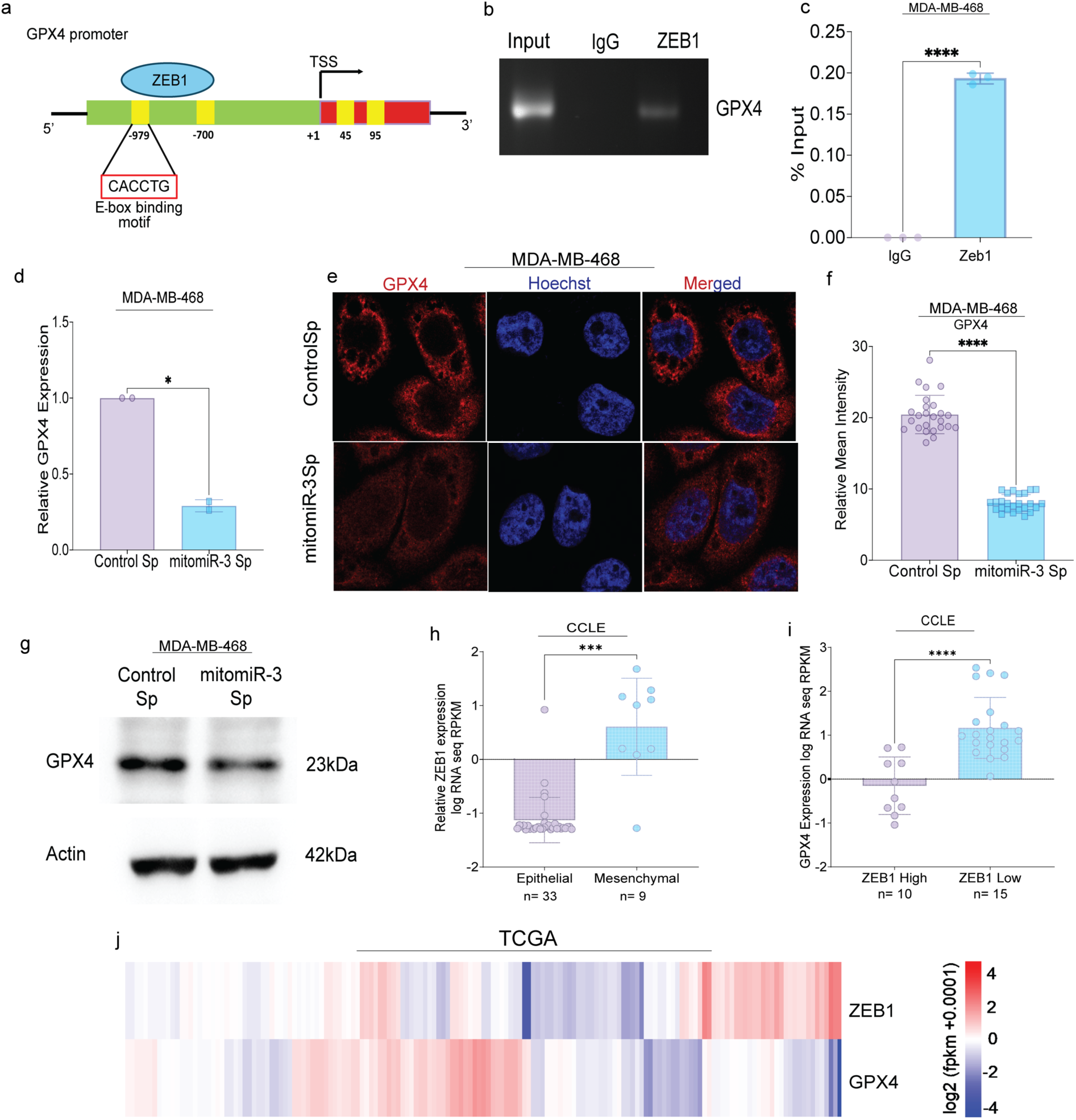
mitomiR-3 inhibition contributes to ZEB1 mediated GPX4 downregulation to promote ferroptosis. a) Schematics of GPX4 promoter sequence, representing E-box location and ZEB1 binding site. b) & c) ChIP-PCR analysis of ZEB1 binding to the GPX4 promoter region in mitomiR-3 Sp transfected MDA-MB-468 cells. RT-PCR was used to detect gene abundance in the different groups, which were immunoprecipitated using anti-ZEB1 antibody in mitomiR-3 Sp cells. The percentage input was calculated by normalizing ZEB1 IP and IgG IP band intensity with input band intensity of the GPX4 RT-PCR product. d) qRT-PCR analysis showing relative GPX4 mRNA expression in mitomiR-3 Sp cells compared to Control Sp cells. Expression of *GPX4* in mitomiR-3 Sp was significantly downregulated when compared to Control Sp. e) Immunofluorescence imaging of GPX4 in Control Sp and mitomiR-3 Sp cells co-stained with Hoechst. Images were acquired using 64x oil immersion objective of confocal microscope (Scale bars, 50μm). A total of 13 different ROIs were analyzed per group. f) Bar graph showing the relative mean intensity of GPX4 in Control Sp and mitomiR-3 Sp cells. mitomiR-3 Sp cells showed reduced GPX4. g) Western blot analysis of GPX4 protein expression in Control Sp and mitomiR-3 Sp cell lines. Densitometric analysis was performed upon normalization of GPX4 protein band intensity to respective β-Actin band. mitomir-3 Sp cells showed decreased protein expression of GPX4; h) Relative ZEB1 expression in epithelial and mesenchymal cell lines (Cancer Cell Line Encyclopedia dataset); i) GPX4 expression in ZEB1 high and ZEB1 low expressing cells (Cancer Cell Line Encyclopedia dataset); j) Heat map showing the relative expression of ZEB1 and GPX4 (MET500). *P-value <0.05, **P-value <0.01, ***P-value <0.001 and ****P-value <0.0001.

### mitomiR-3 sponge promote lipid peroxidation and ferroptosis in TNBC

Lipid peroxidation is the hallmark of ferroptosis ^12,13^. To further confirm the increased lipid peroxidation in mitomiR-3 sponge cells, we used C11 BODIPY 581/591, a fluorescent probe to study lipid peroxidation. Inhibition of mitomiR-3 in TNBC cells significantly increased lipid peroxidation when compared to negative control cells (Fig. 6a, b-c). RSL3 is a known ferroptosis inducer and inhibit glutathione peroxidase gene GPX4 to promote ferroptotic cell death ^32^. We observed increased lipid peroxidation in RSL3 treated TNBC cells (Fig. 6a, d-e). Ferroptosis induction is linked to increased ROS production, lipid peroxidation, defective cellular antioxidant system, perturbed iron metabolism and dysfunctional mitochondrial function.^12,20^ We observed increased cellular ROS, increased lipid peroxidation and decreased total GSH level in mitomiR-3 sponge cells (Fig. 6f-g, Supplementary Fig. 8a-b). Analysis of mitochondrial membrane potential (ΔΨm) and oxygen consumption rate (OCR) demonstrated mitochondrial dysfunction in mitomiR-3 sponge cells when compared to control cells (Supplementary Fig. 8c-h).

**Fig. 6.**
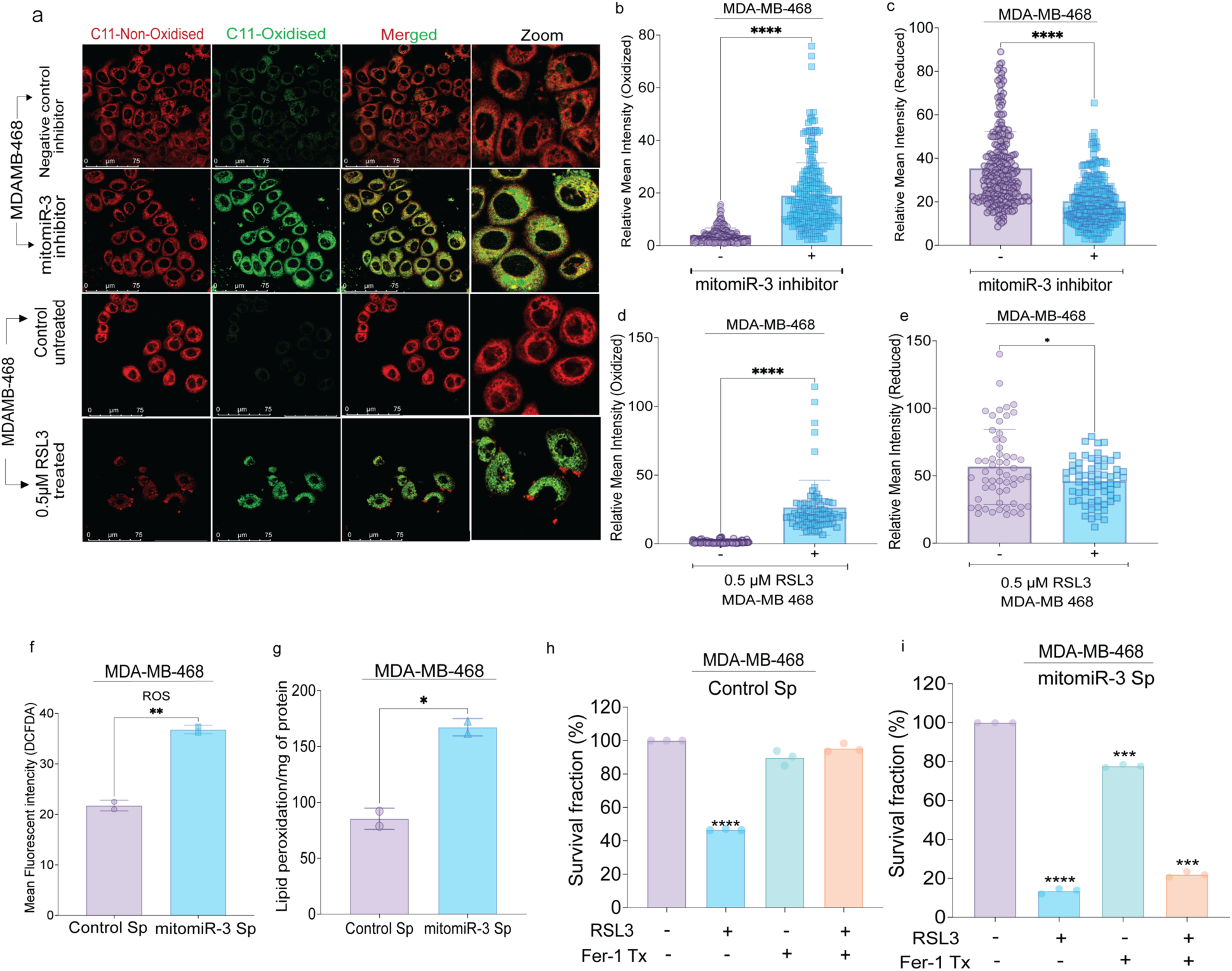
mitomiR-3 regulates lipid ROS and ferroptosis in MDA-MB-468 cells. a) Representative images of the TNBC cell line MDA-MB-468 with 50nM of mitomiR-3 inhibitor and its respective negative control oligonucleotides and MDA-MB-468 treated with 0.5μM RSL3, stained by C11-BODIPY to detect lipid ROS (Scale bar, 75µm). A total of 15 different ROIs were analyzed per group. RSL-3 treated cells were used as a positive control. Increased green fluorescence level of C11-BODIPY was observed in mitomiR-3 Sp cells indicating increased accumulation of lipid ROS triggering lipid peroxidation. b, e) Bar graph showing relative mean intensity of oxidized C11-BODIPY. c, d) Bar graph showing relative mean intensity of reduced C11-BODIPY. f) Bar graph showing flow cytometric analysis of Control Sp and mitomiR-3 Sp cell lines stained for cellular ROS using DCFDA as the molecular probe. Increased accumulation of cellular ROS was observed in mitomiR-3 Sp cells; g) Bar graph showing lipid peroxidation in mitomiR-3 Sp compared to the Control Sp using spectrophotometric LPO assay and normalized to protein concentration of the samples. Lipid peroxidation was significantly increased in mitomiR-3 Sp cells compared to control Sp cells. h, i) mitomiR-3 expression regulates ferroptosis cell death. Bar graph showing mitomiR-3 Sp and Control cells treated with 0.5μM RSL3 and 10μM ferrostatin and cell viability was checked for up to 36hrs. (+) and (−) signs indicate the presence and absence of RSL3 and ferrostatin (Fer-1) respectively. Group with (+) signs for both, were pre-treated with Fer-1, followed by RSL3. On RSL3 treatment control Sp cells showed ∼45% viability and mitomiR-3 cells showed ∼18% viability. In the group that was pre-treatment with ferrostatin-1, followed by RSL3 treatment control Sp cells showed ∼95% viability and mitomiR-3 cells showed ∼20% viability. *p<0.05, **p<0.01, ***p<0.001 and ****p<0.0001.

To understand the impact of mitomiR-3 loss of function on promoting lipid peroxidation and ferroptosis, we treated control (MDA-MB-468) and mitomiR-3 sponge (MDA-MB-468) cells with Fer-1 alone or in combination with RSL3. Ferrostatin-1 inhibits lipid peroxidation and ferroptosis^12,33^. We observed that control cells treated with Fer-1 in combination with RSL3 could rescue the ferroptosis mediated cell death (Fig. 6h). Interestingly, mitomiR-3 sponge cells could not rescue the ferroptosis mediated cell death when treated with Fer-1 in combination with RSL3 (Fig. 6i). Based on the above findings, we could confirm that mitomiR-3 sponge promotes increased lipid peroxidation, mitochondrial dysfunction and ferroptotic cell death in TNBC cells.

## DISCUSSION

Metabolic reprogramming is one of the major hallmarks of cancer cells. Triple-negative breast cancer subtypes are characterized by unique metabolic signatures. As the central hub for multiple metabolic pathways, mitochondria are intricately associated with metabolic reprogramming in TNBC subtypes^34^. Earlier studies characterized TNBC subtypes on the basis of metabolite signatures to identify novel vulnerabilities that can be exploited for developing targeted therapy^35^. In our earlier study, we identified 13 novel mitochondrial genome-encoded miRNAs (mitomiRs) that are differentially expressed in TNBC cells and patient tumors^21^. We observed that mitomiRs are highly expressed in epithelial type TNBC cells (MDA-MB-468) compared with other subtypes. In mesenchymal-type TNBC cells (MDA-MB-231), mitomiRs expression was significantly low which correlated with high ZEB1 expression in MDA-MB-231cells^21^. Here, we uncovered that the majority of the mitomiRs (11 out of 13) have binding sites in the 3′UTR of ZEB1 gene, a core EMT-related transcription factor, in cancer cells. Furthermore, we showed that mitomiR-3, indeed regulates ZEB1 expression in TNBC cells. Expression analysis revealed that the mitomiR-3 level was significantly higher in epithelial-like TNBC cells than in mesenchymal type TNBC cells. Additionally, high mitomiR-3 expression was observed in breast tumor patient tissues. The role of ZEB1 as a major transcriptional regulator of multiple genes involved in cancer cell plasticity, EMT promotion, and regulation of the DNA damage response is well documented^10,22,36^. In our study, we showed that mitomiR-3 loss of function indeed rescued ZEB1 expression in TNBC cells. Multiple studies have shown that EMT is not a binary process in which epithelial cells transform into mesenchymal cells^37^. Rather, it is a gradual process with intermediate cell phenotypes defined as a hybrid EMT state formed by a combination of epithelial and mesenchymal gene expression signatures^38^. The role of miRNAs in regulating the EMT-TFs ZEB1 has been reported^39^. Altered expression of the microRNA-200 family has been observed in cells undergoing EMT^40^. In this study, by manipulating mitomiR-3 expression and function in epithelial-type TNBC cells (MDA-MB-468) and mesenchymal-type TNBC cells (MDA-MB-231) via mimic, inhibitor and sponges, we showed that mitomiR-3 expression (mimic transfected cells) was associated with increased epithelial morphology whereas loss of function (mitomiR-3 inhibitor and sponge transfected cells) showed concomitant increase in mesenchymal morphology. Further, we show that ZEB1 expression in epithelial-type TNBC cells (MDA-MB-468) altered breast cancer cell plasticity, promoting a hybrid EMT phenotype, as confirmed by gene expression analysis, EMT specific protein expression and imaging analysis of cell morphology. Recent studies have shown that mesenchymal-type cancer cells are vulnerable to lipid peroxidation and ferroptosis^17^. Further, it has been shown that enriched PUFA metabolites contribute to sensitivity towards ferroptosis^16,26^. Loss of mitomiR-3 function by miRNA sponge led to metabolic reprogramming in TNBC cells with increased pro-ferroptotic metabolism in TNBC cells. We speculate that mitomiR-3 binds to and regulate the expression of multiple pro-ferroptotic gene expression involved in EMT, iron metabolism, ROS and lipid peroxidation in cancer cells and thereby maintains cell viability. Additionally, we observed increased levels of pro-ferroptotic PUFA metabolites in mitomiR-3 sponge cells suggesting that the mitomiR-3 expression is important for maintaining the level of PUFA metabolites to avoid ferroptosis in TNBC cells. Recent studies have highlighted the roles of the PUFA metabolites arachidonic acid, linolenic acid and dihomo gamma linolenic acid as major contributors to ferroptotic cell death^16,17,19,41,42^. Cancer cells with high ZEB1 expression are vulnerable to ferroptosis but protect themselves by developing a ferroptosis-inhibiting system. We identified that ZEB1 expression in mitomiR-3 sponge cells promoted the transcriptional inhibition of the lipid hydroperoxidase GPX4, contributing to increased ROS mediated mitochondrial dysfunction and lipid peroxidation. These findings are particularly of clinical and translational importance as GPX4 expression contributes to therapy resistance in breast cancer^43^. Interestingly, we observed that the combined enrichment of PUFA metabolites and intracellular iron in mitomiR-3 sponge cells promoted increased lipid peroxidation and ferroptotic cell death. Remarkably, mitomiR-3 sponge cells (loss of function) treated with Ferrostain-1 in combination with RSL3 did not rescue ferroptotic cell death in TNBC cells. Increased mitochondrial ROS in TNBC subtypes promote cell proliferation.^44^ Therefore, we believe that mitomiRs are expressed due to high mitochondrial ROS to inhibit gene expression associated with pro-ferroptotic EMT and lipid peroxidation to avoid ferroptotic cell death in TNBC (Fig. 7). These findings indicated that mitomiR-3 expression in the TNBC cells may be linked to ZEB1 mediated and ZEB1 independent regulation of lipid and iron metabolism contributing to ferroptotic cell death in TNBC. Further, we observed that mitomiR-3 loss of function perturbs TNBC cell proliferation, migration, invasion and *in vivo* tumor growth suggesting mitomiR-3 expression is linked to TNBC cancer cell growth and progression.

**Fig. 7.**
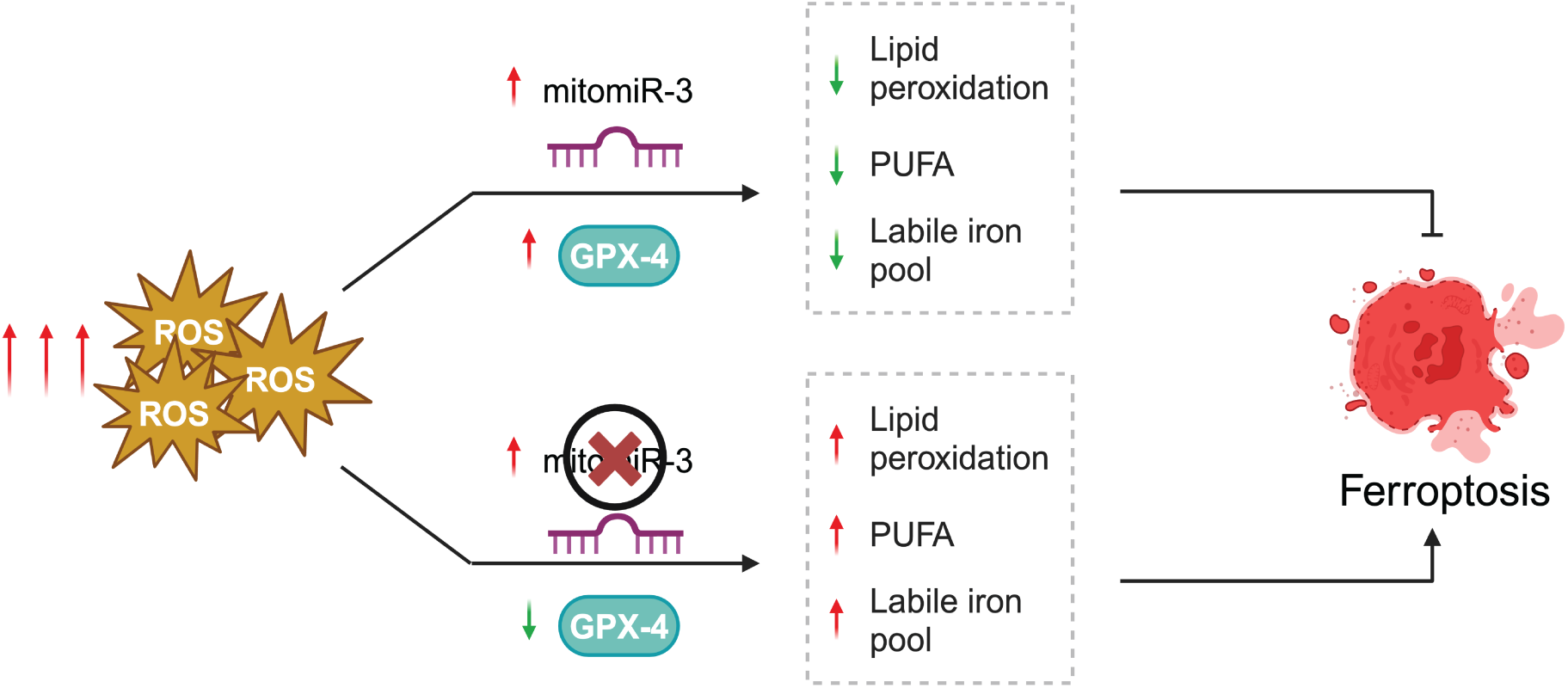
Proposed mechanism of action of mitomiR-3 regulating pro-ferroptotic metabolic signaling in triple-negative breast cancer.

Some of the major drawbacks of the existing ferroptosis inducing drugs include lack of cancer cell specificity, developing resistance, normal tissue toxicity which is severely limiting their translation in clinics. As observed by our study, mitomiRs show high expression in cancer cells and patients which makes them attractive target for induction of ferroptosis. Recent advancements in miRNA based therapeutics have identified multiple strategies to target aberrantly expressed miRNAs (onco-miRs) in cancer cells.^45^ Targeting mitomiR expression in triple-negative breast cancer using anti-mitomiR-3 is an exciting opportunity for developing precise ferroptosis inducing therapy for triple-negative breast cancer patients.

## METHODS

### Cell lines

Breast cancer cell line MCF-7 (Luminal A) and triple negative breast cancer cell lines, MDA-MB-468 (Basal-like TNBC) and MDA-MB-231 (Mesenchymal stem-like TNBC) were cultured in DMEM/F12 (Gibco, USA) supplemented with 10 % Fetal Bovine Serum (FBS) (Gibco, USA). As per standard ATCC recommendation, DMEM/F12 (Gibco, USA) supplemented with 5% Horse serum (Gibco, USA), 0.5μg/μl Hydrocortisone (Himedia), 10μg/μl Insulin (Sigma, USA) and 20pg/μl of Epidermal Growth Factor protein (EGF) (Peprotech, USA) was used to culture MCF10A, immortalized non-malignant human breast epithelial cell line. All cell lines were enzymatically passaged in 0.25% Trypsin-EDTA (HiMedia, India). All cultures were maintained in 37°C incubators at 5% CO2. The culture medium was replenished every two days until cells became 85% confluent.

### Clinical samples

The study was approved by the Kasturba Medical College and Kasturba Hospital Institutional Ethics Committee. The clinical samples were obtained from patients with breast cancer through biopsy at Kasturba Hospital, Manipal, India [(IEC11/2016) and (IEC1:451/2023)] and each patient provided written informed consent. See supplementary Table 1 for more information.

### Animal model

Female athymic BALB/c nude mice (4-6 weeks old) were housed and obtained from the Central Animal Research Facility (Kasturba Medical College, MAHE). All animal experiments were approved by the Research Ethics Committee of Kasturba Medical College, MAHE. All animal handling was performed according to the guidelines for the use and care of live animals by the US National Institutes of Health (NIH).

### Ago2 Immuno-Precipitation (Ago2-IP)

Ago2-IP was performed to investigate the association of mitomiR-3 with Ago2 using previously described protocol ^46^. Briefly, the pure mitoplast fraction and whole cell lysate fractions were lysed in lysis buffer containing Nonidet P-40, RNase inhibitor and protease inhibitor cocktail. The supernatants were collected after centrifugation and were incubated with Ago2 rabbit mAb (1:50) along with equal concentration of rabbit IgG antibody for 6-8 hours at 4°C on a rotospin. Followed by protein: antibody immuno-complex pulldown using Protein G Dynabeads (Invitrogen, USA) overnight at 4°C on a rotospin. The beads were thoroughly washed with wash buffer containing Tween-20 to remove any unbound RNA. RNA was isolated with Trizol reagent for mitomiR and target gene expression ^21^. IgG rabbit antibody was used as an experimental control to normalize for non-specific RNA binding.

### Generation of stable mitomiR-3 loss of function (LoF) sponge cell lines

The loss-of-function model was generated following the protocol previously published.^47^ Briefly, anti-sense mitomiR-3 sponge oligos were cloned into retro-viral expression vector pMSCV-PIG (Puro IRES GFP empty vector) which was a kind gift from David Bartel (Addgene plasmid #21654; http://n2t.net/addgene:21654; RRID: Addgene_21654) ^48^. Initially sense and anti-sense sequences of both linker and sponge sequences were 5’-phosphorylated and mixed in equimolar concentration. This mixture was heated in boiling water for 10 minutes and then slowly cooled to room temperature slowly for 30 minutes to generate oligonucleotide duplex. The duplex was then cloned into the target vector using T4 DNA ligase (NEB, USA) at 1:1000 ratio. Linker sequences were added to introduce Kfl1/SanD1 site into the plasmid between Xho1 and EcoR1 sites. MitomiR-3 sponge oligos were designed with Kfl1/SanD1 sticky ends on both the sides to facilitate directional cloning into linker cloned pMSCV vector. After digestion with Kfl1/SanD1 enzyme, mitomiR-3 sponge duplex were cloned and verified using PCR amplification using plasmid specific primers and sanger DNA sequencing. Clones with multiple copy sponge plasmids was transduced into MDA-MB-468 cell line and selected using 1μg/ml of puromycin (Gibco, USA) antibiotic to generate stable mitomiR-3 sponge models.

### cDNA conversion

Total RNA was isolated from both breast cancer tissue specimens and cell lines. miRNA specific cDNA and total cDNA conversion was performed using TaqMan MicroRNA Reverse Transcription kit and High-Capacity cDNA Reverse Transcription kit (Applied Biosystems, USA) ^21,49^.

### Real time RT-PCR

Real time PCR was performed using custom miRNA TaqMan assays (Applied Biosystems, USA) for mitomiR expression and PowerUp SYBR Green Master Mix (Applied Biosystems, USA) for gene expression analysis. Custom microRNA TaqMan assays were designed as previously described in ^21^. MitomiR expressions were normalized using endogenous control 5S rRNA (for cell line derived RNA) and RNU6B (for tissue derived RNA). For all expression analysis in cell lines, the formula 2^-ΔΔCт^ and for tissue samples the formula 2^-ΔCт^ was used to calculate the relative quantity (RQ).

### Immunoblotting

Cell lysis and immunoblotting was performed as previously described ^21,49^. The cells were lysed using RIPA lysis buffer supplemented with protease inhibitor cocktails and quantified using Bradford method. A total of 30-50μg of protein was resolved on 8-12% SDS-PAGE. Further, proteins were transferred onto 0.45μm nitrocellulose membrane (Bio-Rad, USA) and blocked with 5% non-fat dry milk (Sigma-Aldrich, USA) for respective proteins and incubated for 1 hour at room temperature. The blots were then incubated separately with respective primary antibodies for overnight at 4°C [ZEB1 (1:3000) (Cell Signaling Technologies, USA); GPX4 (1:3000) (ABclonal, Wuhan, China), CDH1 (1:5000) (Cell Signaling Technologies, USA); CDH2 (1:5000) (Cloud Clone, USA); Beta-Actin (1:5000) (Cell Signaling Technologies, USA)]. Horseradish peroxidase (HRP)-conjugated secondary anti-rabbit antibody (1:10000) (Jackson ImmunoResearch Labs, USA) incubation was performed at RT for 2 hours. ECL substrate (BioRad Laboratories, USA) was used to visualize the bands using iBright 1500 (Invitrogen, USA).

### Plasmid construct

ZEB1 3′UTR region with mitomiR-3 binding site was cloned into pmirGLO-Dual Luciferase miRNA target expression vector (Promega, USA) for miRNA luciferase reporter assay (Promega, United States) in MDA-MB-468 cell line. Briefly, ZEB1 3′UTR region was amplified from human genomic DNA, digested using plasmid specific restriction enzymes, and gel eluted along with pmirGLO empty vector. Purified insert and vector DNA were ligated using T4 DNA ligase (NEB, USA) and transformed into DH10β chemical competent cells. Plasmid clones were isolated and verified using restriction digestion. All the clones were confirmed by DNA sequencing before transfecting into MDA-MB-468 cells.

### Transfection and luciferase reporter assay

ZEB1 pmirGLO plasmid along with mitomiR-3 mimic and negative control oligo were co-transfected into MDA-MB-468 cell line using Lipofectamine 3000 reagent (Invitrogen, USA), according to the manufacturer’s instructions. Luciferase assay was performed using Dual-Luciferase Reporter assay (Promega, USA) according to the manufacturer’s instructions using *Renilla* luciferase as internal control after 48 hours.

### Immunofluorescence and Confocal Microscopy

For immunofluorescence, Control Sp and mitomiR-3 Sp cells were seeded on sterile cover slips at 60% confluency. The cells were fixed with 4% paraformaldehyde for 10 min, and permeabilized with Triton X-100 (0.5%) (Sigma-Aldrich, USA). Fixed cells were blocked with 3% BSA (Himedia USA), washed and separately incubated with ZEB1 and GPX4 primary antibodies overnight at 4°C and then labelled with TRITC secondary antibody (Invitrogen, USA). Cells were stained with Hoechst (Invitrogen, USA) for 5min, washed with PBS, and mounted onto a glass slide using Mowiol (Sigma, USA) and stored at 4°C in dark. The images were captured using an SP8-DMi8 confocal microscope (Leica Microsystems, Germany) at 63X magnification ^50^.

### Actin-phalloidin staining

mitomiR-3 mimic, inhibitor, Control Sp and mitomiR-3 Sp transfected cells were grown on coverslips and fixed with 4% paraformaldehyde, stained with ActinRed^TM^ 555 ReadyProbes^TM^ reagent (ThermoFisher Scientific, USA) and Hoechst (Invitrogen, USA), and then mounted onto a glass slide with Mowiol (Sigma, USA) and stored at 4°C in dark. The images were taken at 63X magnification using an SP8-DMi8 confocal microscope (Leica Microsystems, Germany) and processed using Leica Application Suite software (Leica Microsystems, Germany) ^51^.

### RNAseq analysis

Total RNA from Control Sp, mitomiR-3 Sp, and mitomiR-3 mimic transfected MDA-MB- 468 cell lines were isolated and subjected to concentration and quality verification. TruSeq® Stranded mRNA Library Prep (Illumina, Inc.USA) was used for library preparation and mRNA sequencing was performed using NovaSeq 6,000 platform using S4 paired end 200 cycle flow cell according to manufacturer’s protocol. Post sequencing analysis was performed as per the standard Illumina DRAGEN RNA Seq pipelines. Pathway enrichment analysis for the differentially expressed genes (DEGs) from the RNAseq data was performed using Enrichr tool ^52^. The DEGs were clustered into different pathways based on Uniform Manifold Approximation and Projection (UMAP) and ranked based on p-value using Reactome 2022 pathway analysis in Enrichr tool. Gene ontology/KEGG enrichment analysis was performed using Metascape ^53^. Comparative analysis of differentially expressed genes identified in mitomiR-3 sponge cells were performed using DSMZ CellDive web portal.^54^

### Metabolomics analysis

Metabolomic analysis was performed using Control Sp and mitomiR-3 Sp cell supernatant media and the cell pellet. To extract intracellular metabolite pool, cells were washed with chilled PBS and scraped on ice using 200μl of chilled methanol:acetonitrile:water solution [50:30:20(v:v:v)]. Simultaneously, to extract metabolites from the culture media, 200μl of chilled methanol: acetonitrile [50:30 (v:v)] was mixed with 50μl media. Following this both the samples were incubated in −80°C for a minimum of 2-4 hours. To obtain protein-free metabolite extracts, it was subjected to centrifugation twice at 20,000 g for 20 minutes ^55^. The supernatant was vacuum dried, and the pellet was reconstituted using 95:5 ratio of water: acetonitrile with 0.1% formic acid. We then ran the samples under untargeted metabolomics analysis protocol. LC-MS analysis was carried out using Q-Exactive Plus Orbitrap MS (Thermo Scientific, Germany) with a HESI source and Vanquish UHPLC system (Thermo Scientific, Germany). LC–MS/MS data processing was performed using Compound Discoverer 3.1.1 (Thermo Fisher Scientific, USA) software, mainly for peak extraction, peak alignment, and compound identification. Metabolomics data was visualized as a heatmap using SRplot with pheatmap R package ^56^. The peak lists were subjected to Enrichment analysis using the MetaboAnalyst 6.0 (https://www.metaboanalyst.ca/MetaboAnalyst/) online platform ^57^. The parameters used for the enrichment analysis ‘Normalization by median’, log transformation (base 10), Auto scaling, and the KEGG metabolite set library.

### Determination of total ROS

To quantify total cellular ROS, 1 x 10^6^ cells were seeded in 60mm petri plates. Cells were synchronized for 48h, stained with 10μM of DCFH-DA for 30min in the dark at 37°C, and washed with PBS twice for 5min. The pellet was resuspended in PBS and subjected to FACS Calibur flow cytometer (Partec, Germany) and processed using Cell Quest software (BD Biosciences, USA).

### Mitochondrial membrane potential (MMP)

To quantify mitochondrial membrane potential, 1×10^6^ Control Sp and mitomiR-3 Sp cells were plated on 60mm cell culture dish. Cells were serum starved for 48 hours and stained with 100nM of TMRM (Molecular Probes, USA) and incubated for 30 mins at 37°C in dark. Cells will then be washed and resuspended with PBS and subjected to FACS Calibur flow cytometer (Partec, Germany) and processed using Cell Quest software (BD Biosciences, USA).

### Seahorse XFp mitochondrial stress test

To determine the oxygen consumption rate (OCR) in MDA-MB-468 cells transfected with Control Sp and mitomiR-3-3p Sp, Seahorse XFp mitochondrial stress test (Agilent Technologies, USA) was performed according to manufacturer’s protocol. Control Sp and mitomiR-3-3p Sp cells were seeded at a density of 1×10^4^ cells per well in complete media.

The cells were then incubated overnight at 37°C, at 5% CO_2_. Next day, OCR and ECAR values were determined on the Seahorse XFp by running the cell culture miniplates. The media was replaced 1 hour prior to initiation of reading with the test assay medium supplemented with pyruvate, glutamine, and glucose. The injections were in the following order oligomycin, FCCP, and Rotenone/antimycin A. The experiment was performed in triplicates.

### GSH assay

Concentration of GSH was calculated spectrophotometrically by estimating the rate of formation of TNB (5-thio-2-nitrobenzoic acid). Control Sp and mitomiR-3 Sp cells were seeded onto a 100mm cell culture plate at a cell density of 1.5×10^6^. Cells were harvested at 75% confluency and processed with extraction buffer and sonicated. Cells were the centrifuged and the protein estimation was performed by Bradford. Further, 100μl of samples were added per tube and treated with 0.1M KPN and 0.6mM DTNB and incubated at 37°C for 30mins. The reading was spectrophotometrically captured 412nM using an Infinite 200 PRO multimode reader (Tecan, Switzerland) ^58^.

### Lipid peroxidation (LPO)

The thiobarbituric acid assay was used to measure LPO ^59^. Approximately, 1×10^6^cells were seeded in 100mm petri plates and synchronized for 48h. Cells were then lysed using lysis buffer (20mMT ris-HCI pH7.5, 150mM NaCI, ImM EDTA, 1% Triton X-100), centrifuged at 10,000rpm for 10min at 4°C to collect the supernatant. The supernatant was mixed with PBS (pH 7.4) and incubated for 1h at 37°C, followed by Trichloroacetic acid (TCA) addition to a final concentration of 20%. After centrifugation at 10,000rpm for 15min, the supernatant was treated with 1% thiobarbituric acid (TBA), incubated in a boiling water bath for 30min, and the color developed was measured at 532nm using a multimode reader (Tecan, Switzerland).

### RSL3 and Ferrostatin treatment

To understand its role in ferroptosis we treated Control Sp and mitomiR-3 Sp cells with 0.5μM RSL3 (GPX4 mediated ferroptosis inducer) (Sigma-Aldrich, USA) and 10μM ferrostatin-1 (ferroptosis inhibitor) (Sigma-Aldrich, USA) for 3 hours each and then constituted with drug free media for 36 hours. The third group was pre-treated with ferrostatin-1 for 3 hours, followed by treatment with RSL3 for 3 hours and constituted with drug free media for 36 hours. The cell death was estimated by MTT assay at the end of 36 hours.

### BODIPY staining for *in vitro* lipid peroxidation analysis

Control Sp, mitomiR-3-3p Sp and RSL3 treated MDA-MB-468 cells were stained with BODIPY™ 581/591 C11 (Invitrogen, USA) probe, was used as a fluorescent sensor to detect lipid peroxidation using manufacturers protocol. C11-BODIPY’s fluorescence emission peak shifts from[∼[590 nm to[∼[510 nm upon oxidation. Analyses were done by cellular imaging using confocal microscopy. Image quantification was performed using ImageJ. For each cell, the region of interest (ROI) was selected using the Freehand tool in ImageJ, followed by measurement of the mean fluorescence intensity was measured. To eliminate background fluorescence, a small ROI near the selected cell was taken, and its mean fluorescence intensity was measured and recorded. Mean background fluorescence intensity was calculated and Corrected Total Cell Fluorescence (CTCF) for each cell was determined using the formula: CTCF = Integrated Density - (Area of selected cell * Mean background fluorescence intensity).

### ICP-MS detection for iron concentration

ICP-MS (Agilent 7850) was used to estimate the iron load in Control Sp v/s mitomiR-3-3p Sp cells. Cells were seeded at a density of 1×10^6^ in a 60mm cell culture plate and harvested at 75% confluency. The cells were then tested by ICP-MS following standard procedures.

### Chromatin immunoprecipitation (ChIP)

Chromatin immunoprecipitation for mitomiR-3 Sp cells was carried out by crosslinking the cell lysates using 37% formaldehyde. The cells were trypsinized and the extracted chromatin was sonicated for 30 cycles with 30 intervals to get sheared chromatin. The sheared chromatin was incubated with rabbit ZEB1 antibody (1:500) (Cell Signaling Technologies, USA) and equal concentration of rabbit IgG antibodies (1:500) (Cell Signaling Technology, USA) as negative control. 2-5µg of the antibody was added to the diluted chromatin and kept on rotospin overnight at 4°C for efficient binding. The antibody-immunocomplex was captured using Dynabeads protein G (Invitrogen, USA) on a magnetic stand. This complex was subjected to sequential wash with low salt, high salt, LiCl, and TE buffer. The beads were resuspended in an elution buffer and the supernatant containing the sample was collected in a fresh tube. The samples were treated with 5M NaCl, and Proteinase K (200mg/ml) (Sigma-Aldrich, USA), and the phenol-chloroform method was carried out to purify the DNA. Primers for ChIP assay is mentioned in the supplementary table 4. PCR was performed for the target gene.

### Cell proliferation analysis

Control Sp and mitomiR-3 Sp were seeded at a density of 1×10^5^ in 35mm cell culture dishes for up to 72hrs for growth curve analysis. The cells were harvested using trypsin at indicated time points. The cells were stained with trypan blue and the viable cells were counted using a hemocytometer ^51^.

### Apoptotic cell death assay

Annexin-V-FITC/PI staining was performed using control Sp and mitomiR-3 Sp cells to estimate the percentage of apoptotic cells. 1×10^6^ cells were seeded onto 60mm cell culture dish. The percentage of apoptotic cells was quantified by flow cytometry after staining with Annexin-V and PI, as per manufacturer’s instructions (Invitrogen, USA). Cells were then subjected to FACS Calibur flow cytometer (Partec, Germany) and processed using Cell Quest software (BD Biosciences, USA).

### Migration assay

Control Sp and mitomiR-3 Sp cells were cultured to 90% confluency in a 6-well plate. Following PBS wash, cells were cultured in the serum-free medium for 48h. A scratch was made at the center of the plate using a 200μL micro-tip. Subsequently, the cells were cultured in a complete medium, and migration of cells into the scratched area was monitored at the indicated time. Images were captured using Zeiss Primovert microscope equipped with Axiocam 105 camera. Images were processed and analyzed using Zen v2.3 software. The cell migration rate and the wound remaining percentage were estimated as per published protocols ^60^.

### Invasion Assay

*In vitro* cell invasion analysis was performed using the agarose spot assay. In brief, 0.5% agarose solution containing EGF (20 μg/ml) was spotted onto 6-well plates. Control Sp and mitomiR-3 Sp cells (1 × 10^6^) were plated onto wells containing agarose spot in serum-free DMEM medium and incubated at 37 °C for 24h and 48 h. The plates were observed over a period to examine the movement of cells onto the agarose spot. Images of cells were captured every 24hrs. The experiments were performed in duplicates. Images were captured using Zeiss Primovert microscope equipped with Axiocam 105 camera. Images were processed and analyzed using Zen v2.3 software. The cell migration rate and the wound remaining percentage were estimated as per published protocols ^60^

### *In-vivo* Tumorigenicity assay

The Control Sp and mitomiR-3 Sp cells were grafted into 4-6 weeks old female athymic BALB/c nude mice. 3×10^6^ cells/mL were mixed with Matrigel (1:1) (Corning, USA) and subcutaneously injected (n=3 per group) into the lower flank of the nude mice. The tumor growth was monitored for 30 days, and tumor volume measurement was started 10 days post-injection. The tumor dimensions were measured externally with vernier calipers and V = ab2/2 (a, length; b, width) equation was used to calculate the tumor volume. Mice were sacrificed 30 days after injection and tumor tissues were extracted out and images were captured.

### Hematoxylin -Eosin (H&E) staining

Tumor tissue obtained from each animal was fixed in formalin and paraffin blocked were prepared. Tumor tissues were sectioned (5μm) using microtome and stained with H&E (Sigma-Aldrich, USA).

### Immunohistochemistry

Briefly, sections were deparaffinization, antigens were retrieved using a Sodium citrate buffer followed by blocking 1% BSA for 1 hour at room temperature. Sections were separately incubated overnight at 4°C with rabbit anti-ZEB1 (1:500) (Cell Signaling Technologies, USA) and rabbit anti-Ki-67 (1:100) (Invitrogen, USA). Next day the sections were washed with PBST, and blocked with 3% hydrogen peroxide for 15 min, followed by incubating the section with secondary anti-rabbit antibody (1:10000) (Jackson ImmunoResearch Labs, USA) at RT for 2 hours. The sections were covered with DAB substrate (Sigma-Aldrich, USA) for 5 mins and were counterstained with dilute hematoxylin, dehydrated, and mounted for bright-field microscopy.

### TUNEL assay

TUNEL staining was performed using *In Situ* Cell Death Detection Kit, Fluorescein (Roche, Lewes, UK) according to manufacturer’s instructions, to evaluate tissue apoptosis. The percentage apoptotic index for Control Sp and mitomiR-3 Sp cell line-derived xenografts (CDX) was calculated by the formula apoptotic index (%) = 100 x TUNEL positive cells/ total cells.

### Perls’ Prussian blue staining

The tumor samples were stained using a Prussian blue stain. The samples were deparaffinized and subjected to dehydration. Prussian blue staining solution was prepared by mixing potassium ferrocyanide solution and hydrochloric acid in equal proportions. The sections were immersed in the staining solution, washed with distilled water, and subsequently counter-stained with Eosin (Sigma-Aldrich, USA). Finally, the sections subjected to dehydration, then mounted with a neutral mounting medium. The microscopic images were acquired using a microscope ^61^.

### Bioinformatic analysis

*In silico* predictions of mitomiR-3 binding to *ZEB1* was performed using target prediction tools like miRDB ^62^, DIANA Tools ^63^, miRANDA ^64^, RNA22 ^65^, RNAhybrid ^66^ and STarMiR^67^. Multiple sequence alignment (MSA) for seed region conservation analysis of mitomiR-3 across different species was performed using Clustal W ^68^ and visualized in Molecular Evolutionary Genetics Analysis (MEGA 11) software ^69^. Differentially expressed mitomiR-3 target genes in mitomiR-3 mimic and sponge cell datasets were analyzed using public RNA-seq data from 29 breast cancer cell lines in DSMZ CellDive webportal.^54^ To analyze the expression of ZEB1 and GPX4 in metastatic breast cancer, we extracted RNA-seq data from the MET500 cohort in UCSC Xena Browser (https://xena.ucsc.edu/). The mRNA expression levels of ZEB1 and GPX4, measured as log2(FPKM), were visualized in a heatmap across 500 human metastatic cancer samples. For *ZEB1* expression analysis in breast cancer cell lines, mRNA expression data (log RNA seq RPKM) for *ZEB1* were extracted from cBioPortal (http://cbioportal.org). The breast cancer cell lines were shortlisted from the main dataset and classified into epithelial and mesenchymal breast cancer cell lines based on several previously published literature ^70^ and ATCC (https://www.atcc.org/). The mRNA expression of the ZEB1 target gene, GPX4 among ZEB1 high and ZEB1 low breast cancer cell lines was evaluated by extracting mRNA expression (log RNA seq RPKM) for ZEB1 and GPX4 in breast cancer cell lines from cBioPortal (http://cbioportal.org). The breast cancer cells were then classified into ZEB1 high and low breast cancer cells based on their log RNA seq RPKM value. The cell lines with log RNA seq RPKM value above 0 were classified as ZEB1 high breast cancer cell lines and below 0 were classified as ZEB1 low breast cancer cell lines.

### Quantification and statistical analysis

Statistical analysis was performed using GraphPad Prism 10 software. ANOVA and Student t-test (2-tailed unpaired) and data were represented as mean± SD, and a p-value less than 0.05 was considered statistically significant. All the experiments were repeated 2 times and were performed in duplicates.

## ACKNOWLEDGEMENTS

This work was supported by Science and Engineering Research Board (SERB), Department of Science and Technology, Government of India (CRG/2020/004681), Board of Research in Nuclear Sciences (202203HLC19RP07105-BRNS) and Indian Council of Medical Research (ICMR), Government of India (IIRPIG-2023-0000845). RK was supported by ICMR-SRF fellowship (File No: 2019-0278/GEN-BMS), Government of India. The infrastructure support and funding from DST-FIST, TIFAC-CORE, DBT-Builder, VGST Karnataka, K-FIST and Manipal Academy of Higher Education are gratefully acknowledged. We are grateful to the patients for their participation.

## AUTHOR CONTRIBUTIONS

S.C and A.K. conceived and designed the study; A.K. performed experiments with help from J.G., J.S.T, R.K., and R.K.N; A.K., S.M., K.D.M., N.A.K., S.P.K., R.R and Y.S. performed bioinformatic and statistical analysis, generated clinical and experimental data, and compiled figures. A.K. and S.C. wrote the manuscript with input from all other authors. S.C. supervised the study. All authors reviewed and revised the manuscript. S.C. acquired funding for the study.

## DECLARATION OF INTERESTS

The authors declare no competing interest.

## SUPPLEMENTARY MATERIALS

### Supplementary Figure legends

**Supplemental Fig S1**. **Mitochondrial genome encoded miRNAs (mitomiRs) target ZEB1.** (a) Illustration depicting 11 mitomiRs that potentially target ZEB1 predicted using RNA22, RNAhybrid, STarMir and miRanda tools. (b) Schematic representation of predicted binding sites for 11 mitomiRs on the 3’UTR of the ZEB1 mRNA.

**Supplemental Fig S2. Analysis of** mitomiR-3 target gene expression involved in EMT pathway. Heatmap showing expression of EMT pathway genes upregulated in mitomiR-3 sponge cells using public breast cancer cell line RNAseq datasets. TNBC cell lines with mesenchymal features showed high expression of EMT pathway genes upregulated in mitomiR-3 sponge cells.

**Supplemental Fig S3. Inhibition of mitomiR-3 induces the expression of mesenchymal marker gene expression.** (a) Upregulated EMT pathway genes from RNA seq-data in mitomiR-3 Sp and Control Sp. Enrichr tool was used to plot the bokeh graph showing the gene enrichment pattern, clustered based on Uniform Manifold Approximation and Projection (UMAP). (b) Bar graph of GO/KEGG enrichment analysis using Metascape.

**Supplemental Fig S4. mitomiR-3 loss of function promotes mesenchymal phenotype in mitomiR-3 inhibitor treated and mitomiR-3 sponge TNBC cells.** (a) Bright field microscopy (20x magnification) of the TNBC cell lines MDA-MB-231 transfected with 50nM of mitomiR-3 mimic and its respective negative control oligonucleotides. The bar graph shows the percentage of epithelial cells. MDA-MB-231 (MSL) cells transfected with mitomiR-3 mimic showed an increase in the number of cells showing epithelial phenotype. (b) Bright field microscopy (20x magnification) imaging of the TNBC cell lines MDA-MB- 468 transfected with 50nM of mitomiR-3 inhibitor and its respective negative control oligonucleotides. Bar graph showing the percentage of mesenchymal cells. MDA-MB-468 (BL-1) cells transfected with mitomiR-3 inhibitor showed an increase in the number of cells showing mesenchymal phenotype. (c) Representative images of actin phalloidin staining in mitomiR-3 Sp and Control Sp (Scale bars, 50µm). A total of 12 different ROIs were analyzed per group. The bar graph shows an increase in cellular area, cellular perimeter and aspect ratio in mitomiR-3 Sp cells. mitomiR-3 Sp cells showed increased mesenchymal cell type specific features.

**Supplemental Fig S5. Inhibition of mitomiR-3 increased cell death in mitomiR-3 sponge TNBC cells.** Annexin V-FITC apoptosis assay to assess difference between mitomiR-3 Sp and Control Sp. *p<0.05, **p<0.01, ***p<0.001 and ****p<0.0001.

**Supplemental Fig S6. Pathway analysis of metabolomics data showed enrichment of PUFA metabolism in mitomiR-3 sponge TNBC cells.** The bubble plot represents the metabolite set enrichment in mitomiR-3 Sp cells. The plot showed high enrichment for lipid metabolism pathways.

**Supplemental Fig S7. Inhibition of mitomiR-3 reprograms lipid and iron metabolism in mitomiR-3 sponge TNBC cells.** a) Pathway analysis of upregulated genes in mitomiR-3 LOF cells showed enrichment in lipid metabolism. b) Heatmap showing the expression (log FC) of genes associated with iron metabolism in mitomiR-3 Sp and mitomiR-3 mimic. c) Bar plot showing iron content (ppb) in Control Sp and mitomiR-3 Sp. mitomiR-3 Sp had increased accumulation of iron when compared to control Sp. *p<0.05.

**Supplemental Fig S8. Inhibition of mitomiR-3 reduces cellular glutathione, mitochondrial membrane potential and oxygen consumption rate in mitomiR-3 sponge TNBC cells.** (a) The graph represents intracellular glutathione levels quantified using spectrophotometric glutathione assay in Control Sp and mitomiR-3 Sp. Intracellular GSH is significantly reduced in mitomiR-3 Sp cells compare to the Control Sp cells; (b) Flow cytometry histogram illustrating ROS content by DCFDA staining. (c) Bar graph showing flow cytometric analysis of Control Sp and mitomiR-3 Sp cell lines stained for mitochondrial membrane potential (TMRM). The mitochondrial membrane potential was significantly reduced in mitomiR-3 Sp cells. (d) Flow cytometry histogram illustrating mitochondrial membrane potential by TMRM staining; (e) Representative graph showing the OCR in Control Sp and mitomiR-3 Sp cell lines. Injections of oligomycin, FCCP, and antimycin A and rotenone (A/R) are indicated. (f–j) Representative bar graph quantifying different mitochondrial bioenergetic parameters: maximal respiration, nonmitochondrial oxygen consumption, proton leak, spare respiratory capacity and spare respiratory capacity (%). *p<0.05, **p<0.01.

## Notes

### Competing Interest Statement

The authors have declared no competing interest.

## REFERENCES

1. Lehmann, B.D., Bauer, J.A., Chen, X., Sanders, M.E., Chakravarthy, A.B., Shyr, Y., and Pietenpol, J.A. (2011). Identification of human triple-negative breast cancer subtypes and preclinical models for selection of targeted therapies. Journal of Clinical Investigation 121, 2750–2767. 10.1172/JCI45014.

2. Lehmann, B.D., and Pietenpol, J.A. (2014). Identification and use of biomarkers in treatment strategies for triple-negative breast cancer subtypes. J Pathol 232, 142–150. 10.1002/path.4280.

3. Lehmann, B.D., Jovanović, B., Chen, X., Estrada, M.V., Johnson, K.N., Shyr, Y., Moses, H.L., Sanders, M.E., and Pietenpol, J.A. (2016). Refinement of Triple-Negative Breast Cancer Molecular Subtypes: Implications for Neoadjuvant Chemotherapy Selection. PLoS One 11, e0157368. 10.1371/journal.pone.0157368.

4. Marra, A., Trapani, D., Viale, G., Criscitiello, C., and Curigliano, G. (2020). Practical classification of triple-negative breast cancer: intratumoral heterogeneity, mechanisms of drug resistance, and novel therapies. NPJ Breast Cancer 6, 54. 10.1038/s41523-020-00197-2.

5. Yin, L., Duan, J.J., Bian, X.W., and Yu, S.C. (2020). Triple-negative breast cancer molecular subtyping and treatment progress. Breast Cancer Research 22, 1–13. 10.1186/s13058-020-01296-5.

6. Mohammadi Ghahhari, N., Sznurkowska, M.K., Hulo, N., Bernasconi, L., Aceto, N., and Picard, D. (2022). Cooperative interaction between ERα and the EMT-inducer ZEB1 reprograms breast cancer cells for bone metastasis. Nat Commun 13, 2104. 10.1038/s41467-022-29723-5.

7. Yang, J., Antin, P., Berx, G., Blanpain, C., Brabletz, T., Bronner, M., Campbell, K., Cano, A., Casanova, J., Christofori, G., et al. (2020). Guidelines and definitions for research on epithelial-mesenchymal transition. Nat Rev Mol Cell Biol 21, 341–352. 10.1038/s41580-020-0237-9.

8. Taube, J.H., Herschkowitz, J.I., Komurov, K., Zhou, A.Y., Gupta, S., Yang, J., Hartwell, K., Onder, T.T., Gupta, P.B., Evans, K.W., et al. (2010). Core epithelial-to-mesenchymal transition interactome gene-expression signature is associated with claudin-low and metaplastic breast cancer subtypes. Proceedings of the National Academy of Sciences 107, 15449–15454. 10.1073/pnas.1004900107.

9. Chaffer, C.L., Marjanovic, N.D., Lee, T., Bell, G., Kleer, C.G., Reinhardt, F., D’Alessio, A.C., Young, R.A., and Weinberg, R.A. (2013). Poised chromatin at the ZEB1 promoter enables breast cancer cell plasticity and enhances tumorigenicity. Cell 154, 61–74. 10.1016/j.cell.2013.06.005.

10. Krebs, A.M., Mitschke, J., Lasierra Losada, M., Schmalhofer, O., Boerries, M., Busch, H., Boettcher, M., Mougiakakos, D., Reichardt, W., Bronsert, P., et al. (2017). The EMT-activator Zeb1 is a key factor for cell plasticity and promotes metastasis in pancreatic cancer. Nat Cell Biol 19, 518–529. 10.1038/ncb3513.

11. Wu, H.-T., Zhong, H.-T., Li, G.-W., Shen, J.-X., Ye, Q.-Q., Zhang, M.-L., and Liu, J. (2020). Oncogenic functions of the EMT-related transcription factor ZEB1 in breast cancer. J Transl Med 18, 51. 10.1186/s12967-020-02240-z.

12. Dixon, S.J., Lemberg, K.M., Lamprecht, M.R., Skouta, R., Zaitsev, E.M., Gleason, C.E., Patel, D.N., Bauer, A.J., Cantley, A.M., Yang, W.S., et al. (2012). Ferroptosis: An Iron-Dependent Form of Nonapoptotic Cell Death. Cell 149, 1060–1072. 10.1016/j.cell.2012.03.042.

13. Yang, W.S., SriRamaratnam, R., Welsch, M.E., Shimada, K., Skouta, R., Viswanathan, V.S., Cheah, J.H., Clemons, P.A., Shamji, A.F., Clish, C.B., et al. (2014). Regulation of ferroptotic cancer cell death by GPX4. Cell 156, 317–331. 10.1016/j.cell.2013.12.010.

14. Schwab, A., Rao, Z., Zhang, J., Gollowitzer, A., Siebenkäs, K., Bindel, N., D’Avanzo, E., Van Roey, R., Hajjaj, Y., Özel, E., et al. (2024). Zeb1 mediates EMT/plasticity-associated ferroptosis sensitivity in cancer cells by regulating lipogenic enzyme expression and phospholipid composition. Nat Cell Biol. 10.1038/s41556-024-01464-1.

15. Doll, S., Proneth, B., Tyurina, Y.Y., Panzilius, E., Kobayashi, S., Ingold, I., Irmler, M., Beckers, J., Aichler, M., Walch, A., et al. (2017). ACSL4 dictates ferroptosis sensitivity by shaping cellular lipid composition. Nat Chem Biol 13, 91–98. 10.1038/nchembio.2239.

16. Qiu, B., Zandkarimi, F., Bezjian, C.T., Reznik, E., Soni, R.K., Gu, W., Jiang, X., and Stockwell, B.R. (2024). Phospholipids with two polyunsaturated fatty acyl tails promote ferroptosis. Cell 187, 1177–1190.e18. 10.1016/j.cell.2024.01.030.

17. Lee, J.-Y., Nam, M., Son, H.Y., Hyun, K., Jang, S.Y., Kim, J.W., Kim, M.W., Jung, Y., Jang, E., Yoon, S.-J., et al. (2020). Polyunsaturated fatty acid biosynthesis pathway determines ferroptosis sensitivity in gastric cancer. Proceedings of the National Academy of Sciences 117, 32433–32442. 10.1073/pnas.2006828117.

18. Suda, A., Umaru, B.A., Yamamoto, Y., Shima, H., Saiki, Y., Pan, Y., Jin, L., Sun, J., Low, Y.L.C., Suzuki, C., et al. (2024). Polyunsaturated fatty acids-induced ferroptosis suppresses pancreatic cancer growth. Sci Rep 14, 4409. 10.1038/s41598-024-55050-4.

19. Lorito, N., Subbiani, A., Smiriglia, A., Bacci, M., Bonechi, F., Tronci, L., Romano, E., Corrado, A., Longo, D.L., Iozzo, M., et al. (2024). FADS1/2 control lipid metabolism and ferroptosis susceptibility in triple-negative breast cancer. EMBO Mol Med 16, 1533– 1559. 10.1038/s44321-024-00090-6.

20. Gao, M., Yi, J., Zhu, J., Minikes, A.M., Monian, P., Thompson, C.B., and Jiang, X. (2019). Role of Mitochondria in Ferroptosis. Mol Cell 73, 354–363.e3. 10.1016/j.molcel.2018.10.042.

21. Kuthethur, R., Shukla, V., Mallya, S., Adiga, D., Kabekkodu, S.P., Ramachandra, L., Saxena, P.U.P., Satyamoorthy, K., and Chakrabarty, S. (2022). Expression analysis and function of mitochondrial genome-encoded microRNAs. J Cell Sci 135, jcs258937. 10.1242/jcs.258937.

22. Aghdassi, A., Sendler, M., Guenther, A., Mayerle, J., Behn, C.-O., Heidecke, C.-D., Friess, H., Büchler, M., Evert, M., Lerch, M.M., et al. (2012). Recruitment of histone deacetylases HDAC1 and HDAC2 by the transcriptional repressor ZEB1 downregulates E-cadherin expression in pancreatic cancer. Gut 61, 439–448. 10.1136/gutjnl-2011-300060.

23. Ahn, Y.-H., Gibbons, D.L., Chakravarti, D., Creighton, C.J., Rizvi, Z.H., Adams, H.P., Pertsemlidis, A., Gregory, P.A., Wright, J.A., Goodall, G.J., et al. (2012). ZEB1 drives prometastatic actin cytoskeletal remodeling by downregulating miR-34a expression. J Clin Invest 122, 3170–3183. 10.1172/JCI63608.

24. Seiler, A., Schneider, M., Förster, H., Roth, S., Wirth, E.K., Culmsee, C., Plesnila, N., Kremmer, E., Rådmark, O., Wurst, W., et al. (2008). Glutathione peroxidase 4 senses and translates oxidative stress into 12/15-lipoxygenase dependent- and AIF-mediated cell death. Cell Metab 8, 237–248. 10.1016/j.cmet.2008.07.005.

25. Singh, N.K., and Rao, G.N. (2019). Emerging role of 12/15-Lipoxygenase (ALOX15) in human pathologies. Prog Lipid Res 73, 28–45. 10.1016/j.plipres.2018.11.001.

26. Kagan, V.E., Mao, G., Qu, F., Angeli, J.P.F., Doll, S., Croix, C.S., Dar, H.H., Liu, B., Tyurin, V.A., Ritov, V.B., et al. (2017). Oxidized arachidonic and adrenic PEs navigate cells to ferroptosis. Nat Chem Biol 13, 81–90. 10.1038/nchembio.2238.

27. Li, Y., Du, Y., Zhou, Y., Chen, Q., Luo, Z., Ren, Y., Chen, X., and Chen, G. (2023). Iron and copper: critical executioners of ferroptosis, cuproptosis and other forms of cell death. Cell Communication and Signaling 21, 327. 10.1186/s12964-023-01267-1.

28. Shen, Z., Liu, T., Li, Y., Lau, J., Yang, Z., Fan, W., Zhou, Z., Shi, C., Ke, C., Bregadze, V.I., et al. (2018). Fenton-Reaction-Acceleratable Magnetic Nanoparticles for Ferroptosis Therapy of Orthotopic Brain Tumors. ACS Nano 12, 11355–11365. 10.1021/acsnano.8b06201.

29. Yan, H., Zou, T., Tuo, Q., Xu, S., Li, H., Belaidi, A.A., and Lei, P. (2021). Ferroptosis: mechanisms and links with diseases. Sig Transduct Target Ther 6, 1–16. 10.1038/s41392-020-00428-9.

30. Barretina, J., Caponigro, G., Stransky, N., Venkatesan, K., Margolin, A.A., Kim, S., Wilson, C.J., Lehár, J., Kryukov, G.V., Sonkin, D., et al. (2012). The Cancer Cell Line Encyclopedia enables predictive modelling of anticancer drug sensitivity. Nature 483, 603–607. 10.1038/nature11003.

31. Robinson, D.R., Wu, Y.-M., Lonigro, R.J., Vats, P., Cobain, E., Everett, J., Cao, X., Rabban, E., Kumar-Sinha, C., Raymond, V., et al. (2017). Integrative clinical genomics of metastatic cancer. Nature 548, 297–303. 10.1038/nature23306.

32. Yang, W.S., and Stockwell, B.R. (2008). Synthetic lethal screening identifies compounds activating iron-dependent, nonapoptotic cell death in oncogenic-RAS-harboring cancer cells. Chem Biol 15, 234–245. 10.1016/j.chembiol.2008.02.010.

33. Skouta, R., Dixon, S.J., Wang, J., Dunn, D.E., Orman, M., Shimada, K., Rosenberg, P.A., Lo, D.C., Weinberg, J.M., Linkermann, A., et al. (2014). Ferrostatins Inhibit Oxidative Lipid Damage and Cell Death in Diverse Disease Models. J. Am. Chem. Soc. 136, 4551–4556. 10.1021/ja411006a.

34. Pelicano, H., Zhang, W., Liu, J., Hammoudi, N., Dai, J., Xu, R.-H., Pusztai, L., and Huang, P. (2014). Mitochondrial dysfunction in some triple-negative breast cancer cell lines: role of mTOR pathway and therapeutic potential. Breast Cancer Research 16, 434. 10.1186/s13058-014-0434-6.

35. Yang, F., Xiao, Y., Ding, J.-H., Jin, X., Ma, D., Li, D.-Q., Shi, J.-X., Huang, W., Wang, Y.-P., Jiang, Y.-Z., et al. (2023). Ferroptosis heterogeneity in triple-negative breast cancer reveals an innovative immunotherapy combination strategy. Cell Metabolism 35, 84-100.e8. 10.1016/j.cmet.2022.09.021.

36. Genetta, T.L., Hurwitz, J.C., Clark, E.A., Herold, B.T., Khalil, S., Abbas, T., and Larner, J.M. (2023). ZEB1 promotes non-homologous end joining double-strand break repair. Nucleic Acids Research 51, 9863–9879. 10.1093/nar/gkad723.

37. Kröger, C., Afeyan, A., Mraz, J., Eaton, E.N., Reinhardt, F., Khodor, Y.L., Thiru, P., Bierie, B., Ye, X., Burge, C.B., et al. (2019). Acquisition of a hybrid E/M state is essential for tumorigenicity of basal breast cancer cells. Proceedings of the National Academy of Sciences 116, 7353–7362. 10.1073/pnas.1812876116.

38. Schliekelman, M.J., Taguchi, A., Zhu, J., Dai, X., Rodriguez, J., Celiktas, M., Zhang, Q., Chin, A., Wong, C.-H., Wang, H., et al. (2015). Molecular portraits of epithelial, mesenchymal and hybrid states in lung adenocarcinoma and their relevance to survival. Cancer Res 75, 1789–1800. 10.1158/0008-5472.CAN-14-2535.

39. Bracken, C.P., Gregory, P.A., Kolesnikoff, N., Bert, A.G., Wang, J., Shannon, M.F., and Goodall, G.J. (2008). A double-negative feedback loop between ZEB1-SIP1 and the microRNA-200 family regulates epithelial-mesenchymal transition. Cancer Res 68, 7846– 7854. 10.1158/0008-5472.CAN-08-1942.

40. Gregory, P.A., Bert, A.G., Paterson, E.L., Barry, S.C., Tsykin, A., Farshid, G., Vadas, M.A., Khew-Goodall, Y., and Goodall, G.J. (2008). The miR-200 family and miR-205 regulate epithelial to mesenchymal transition by targeting ZEB1 and SIP1. Nat Cell Biol 10, 593– 601. 10.1038/ncb1722.

41. Beatty, A., Singh, T., Tyurina, Y.Y., Tyurin, V.A., Samovich, S., Nicolas, E., Maslar, K., Zhou, Y., Cai, K.Q., Tan, Y., et al. (2021). Ferroptotic cell death triggered by conjugated linolenic acids is mediated by ACSL1. Nat Commun 12, 2244. 10.1038/s41467-021-22471-y.

42. Sarparast, M., Pourmand, E., Hinman, J., Vonarx, D., Reason, T., Zhang, F., Paithankar, S., Chen, B., Borhan, B., Watts, J.L., et al. (2023). Dihydroxy-Metabolites of Dihomo-γ-linolenic Acid Drive Ferroptosis-Mediated Neurodegeneration. ACS Cent Sci 9, 870–882. 10.1021/acscentsci.3c00052.

43. Hangauer, M.J., Viswanathan, V.S., Ryan, M.J., Bole, D., Eaton, J.K., Matov, A., Galeas, J., Dhruv, H.D., Berens, M.E., Schreiber, S.L., et al. (2017). Drug-tolerant persister cancer cells are vulnerable to GPX4 inhibition. Nature 551, 247–250. 10.1038/nature24297.

44. Sarmiento-Salinas, F.L., Delgado-Magallón, A., Montes-Alvarado, J.B., Ramírez-Ramírez, D., Flores-Alonso, J.C., Cortés-Hernández, P., Reyes-Leyva, J., Herrera-Camacho, I., Anaya-Ruiz, M., Pelayo, R., et al. (2019). Breast Cancer Subtypes Present a Differential Production of Reactive Oxygen Species (ROS) and Susceptibility to Antioxidant Treatment. Frontiers in Oncology 9, 480. 10.3389/fonc.2019.00480.

45. Wang, Y., Malik, S., Suh, H.-W., Xiao, Y., Deng, Y., Fan, R., Huttner, A., Bindra, R.S., Singh, V., Saltzman, W.M., et al. (2023). Anti-seed PNAs targeting multiple oncomiRs for brain tumor therapy. Science Advances 9, eabq7459. 10.1126/sciadv.abq7459.

46. Tan, L.P., Seinen, E., Duns, G., de Jong, D., Sibon, O.C.M., Poppema, S., Kroesen, B.-J., Kok, K., and van den Berg, A. (2009). A high throughput experimental approach to identify miRNA targets in human cells. Nucleic Acids Res 37, e137. 10.1093/nar/gkp715.

47. Kluiver, J., Slezak-Prochazka, I., Smigielska-Czepiel, K., Halsema, N., Kroesen, B.-J., and van den Berg, A. (2012). Generation of miRNA sponge constructs. Methods 58, 113–117. 10.1016/j.ymeth.2012.07.019.

48. Mayr, C., and Bartel, D.P. (2009). Widespread shortening of 3’UTRs by alternative cleavage and polyadenylation activates oncogenes in cancer cells. Cell 138, 673–684. 10.1016/j.cell.2009.06.016.

49. Kuthethur, R., Adiga, D., Kandettu, A., Jerome, M.S., Mallya, S., Mumbrekar, K.D., Kabekkodu, S.P., and Chakrabarty, S. (2023). MiR-4521 perturbs FOXM1-mediated DNA damage response in breast cancer. Front Mol Biosci 10, 1131433. 10.3389/fmolb.2023.1131433.

50. Bhat, S., Adiga, D., Shukla, V., Guruprasad, K.P., Kabekkodu, S.P., and Satyamoorthy, K. (2022). Metastatic suppression by DOC2B is mediated by inhibition of epithelial-mesenchymal transition and induction of senescence. Cell Biol Toxicol 38, 237–258. 10.1007/s10565-021-09598-w.

51. Bhat, S., Kabekkodu, S.P., Adiga, D., Fernandes, R., Shukla, V., Bhandari, P., Pandey, D., Sharan, K., and Satyamoorthy, K. (2021). ZNF471 modulates EMT and functions as methylation regulated tumor suppressor with diagnostic and prognostic significance in cervical cancer. Cell Biol Toxicol 37, 731–749. 10.1007/s10565-021-09582-4.

52. Kuleshov, M.V., Jones, M.R., Rouillard, A.D., Fernandez, N.F., Duan, Q., Wang, Z., Koplev, S., Jenkins, S.L., Jagodnik, K.M., Lachmann, A., et al. (2016). Enrichr: a comprehensive gene set enrichment analysis web server 2016 update. Nucleic Acids Res 44, W90–97. 10.1093/nar/gkw377.

53. Zhou, Y., Zhou, B., Pache, L., Chang, M., Khodabakhshi, A.H., Tanaseichuk, O., Benner, C., and Chanda, S.K. (2019). Metascape provides a biologist-oriented resource for the analysis of systems-level datasets. Nat Commun 10, 1523. 10.1038/s41467-019-09234-6.

54. Koblitz, J., Dirks, W.G., Eberth, S., Nagel, S., Steenpass, L., and Pommerenke, C. (2022). DSMZCellDive: Diving into high-throughput cell line data. F1000Res 11, 420. 10.12688/f1000research.111175.2.

55. Ahn, E., Kumar, P., Mukha, D., Tzur, A., and Shlomi, T. (2017). Temporal fluxomics reveals oscillations in TCA cycle flux throughout the mammalian cell cycle. Mol Syst Biol 13, 953. 10.15252/msb.20177763.

56. Tang, D., Chen, M., Huang, X., Zhang, G., Zeng, L., Zhang, G., Wu, S., and Wang, Y. (2023). SRplot: A free online platform for data visualization and graphing. PLoS One 18, e0294236. 10.1371/journal.pone.0294236.

57. Pang, Z., Lu, Y., Zhou, G., Hui, F., Xu, L., Viau, C., Spigelman, A.F., MacDonald, P.E., Wishart, D.S., Li, S., et al. (2024). MetaboAnalyst 6.0: towards a unified platform for metabolomics data processing, analysis and interpretation. Nucleic Acids Res, gkae253. 10.1093/nar/gkae253.

58. Rahman, I., Kode, A., and Biswas, S.K. (2006). Assay for quantitative determination of glutathione and glutathione disulfide levels using enzymatic recycling method. Nat Protoc 1, 3159–3165. 10.1038/nprot.2006.378.

59. Garcia, Y.J., Rodríguez-Malaver, A.J., and Peñaloza, N. (2005). Lipid peroxidation measurement by thiobarbituric acid assay in rat cerebellar slices. J Neurosci Methods 144, 127–135. 10.1016/j.jneumeth.2004.10.018.

60. Xu, N., Zhang, L., Meisgen, F., Harada, M., Heilborn, J., Homey, B., Grandér, D., Ståhle, M., Sonkoly, E., and Pivarcsi, A. (2012). MicroRNA-125b down-regulates matrix metallopeptidase 13 and inhibits cutaneous squamous cell carcinoma cell proliferation, migration, and invasion. J Biol Chem 287, 29899–29908. 10.1074/jbc.M112.391243.

61. Ghio, A.J., and Roggli, V.L. (2021). Perls’ Prussian Blue Stains of Lung Tissue, Bronchoalveolar Lavage, and Sputum. J Environ Pathol Toxicol Oncol 40, 1–15. 10.1615/JEnvironPatholToxicolOncol.2020036292.

62. Chen, Y., and Wang, X. (2020). miRDB: an online database for prediction of functional microRNA targets. Nucleic Acids Res 48, D127–D131. 10.1093/nar/gkz757.

63. Vlachos, I.S., and Hatzigeorgiou, A.G. (2017). Functional Analysis of miRNAs Using the DIANA Tools Online Suite. Methods Mol Biol 1517, 25–50. 10.1007/978-1-4939-6563-2_2.

64. Enright, A.J., John, B., Gaul, U., Tuschl, T., Sander, C., and Marks, D.S. (2003). MicroRNA targets in Drosophila. Genome Biol 5, R1. 10.1186/gb-2003-5-1-r1.

65. Miranda, K.C., Huynh, T., Tay, Y., Ang, Y.-S., Tam, W.-L., Thomson, A.M., Lim, B., and Rigoutsos, I. (2006). A pattern-based method for the identification of MicroRNA binding sites and their corresponding heteroduplexes. Cell 126, 1203–1217. 10.1016/j.cell.2006.07.031.

66. Rehmsmeier, M., Steffen, P., Hochsmann, M., and Giegerich, R. (2004). Fast and effective prediction of microRNA/target duplexes. RNA 10, 1507–1517. 10.1261/rna.5248604.

67. Rennie, W., Liu, C., Carmack, C.S., Wolenc, A., Kanoria, S., Lu, J., Long, D., and Ding, Y. (2014). STarMir: a web server for prediction of microRNA binding sites. Nucleic Acids Res 42, W114–118. 10.1093/nar/gku376.

68. Thompson, J.D., Gibson, T.J., and Higgins, D.G. (2002). Multiple sequence alignment using ClustalW and ClustalX. Curr Protoc Bioinformatics Chapter 2, Unit 2.3. 10.1002/0471250953.bi0203s00.

69. Tamura, K., Stecher, G., and Kumar, S. (2021). MEGA11: Molecular Evolutionary Genetics Analysis Version 11. Mol Biol Evol 38, 3022–3027. 10.1093/molbev/msab120.

70. Chakraborty, P., George, J.T., Tripathi, S., Levine, H., and Jolly, M.K. (2020). Comparative Study of Transcriptomics-Based Scoring Metrics for the Epithelial-Hybrid-Mesenchymal Spectrum. Front Bioeng Biotechnol 8, 220. 10.3389/fbioe.2020.00220.

