## Supplemental tables for "Mitochondrial genome-encoded mitomiRs regulate cellular plasticity and susceptibility to ferroptosis in triple-negative breast cancer"

**SUPPLEMENTAL INFORMATION**

**Supplementary Table 1:** Clinical data of the breast cancer patient samples used for analysis of mitomiR-3 expression.

| **Breast Cancer Patient Clinicopathological Details** | |
| --- | --- |
| **Number of patients** | 15 |
| **Age** | |
| **<50** | 7 |
| **>50** | 8 |
| **Intrinsic subtypes** | |
| **Luminal A (ER+, PR+/-, Her2-)** | 4 |
| **Luminal B (ER+, PR+/-, Her2+)** | 4 |
| **Triple Negative (ER-, PR-, Her2-)** | 6 |
| **Her2 Positive (ER-, PR-, Her2+)** | 1 |
| **Tumor grade** | |
| **Grade 1** | 1 |
| **Grade 2** | 5 |
| **Grade 3** | 9 |
| **Lymph Node Status** | |
| **Positive** | 11 |
| **Negative** | 4 |

**Supplementary Table 2**: Expression analysis of mitomiR-3 target genes involved in epithelial to mesenchymal pathway (EMT) genes in mitomiR-3 mimic (gain of function) transfected and mitomiR-3 sponge (loss of function) transfected TNBC cells.

| **Gene Symbol mimic** | **mitomiR-3 mimic logFC** | **mitomiR-3 Target Gene** | **mitomiR-3 Sponge logFC** | **Description** | **Reference** |
| --- | --- | --- | --- | --- | --- |
| VIM | -0.64 | Yes | 0.94 | Vimentin | ^1^ |
| LCN2 | -1.47 | Yes | 2.76 | Lipocalin-2 | ^2^ |
| CAV1 | -0.58 | Yes | 2.27 | Caveolin-1 | ^3^ |
| LOX | -0.42 | Yes | 1.02 | Lysyl oxidase | ^4^ |
| MET | -0.31 | Yes | 1.72 | MET proto-oncogene, receptor tyrosine kinase | ^5^ |
| CLDN1 | -0.52 | Yes | 0.73 | Claudin-1 | ^6^ |
| SIRT1 | -0.47 | Yes | 0.33 | Sirtuin-1 | ^7^ |
| FOSL1 | -0.31 | Yes | 0.41 | FOS Like 1 | ^8^ |
| MCAM | -0.33 | Yes | 2.59 | Melanoma Cell Adhesion Molecule | ^9^ |
| ZEB1 | -0.23 | Yes | 3.87 | Zinc Finger E-Box Binding Homeobox 1 | ^10^ |
| MACC1 | -2.59 | Yes | 0.94 | Metastasis Associated In Colon Cancer 1 | ^11^ |
| DAB2 | -0.25 | Yes | 1.11 | Disabled-2 | ^12^ |
| EGFR | -0.11 | Yes | 0.63 | Epidermal Growth Factor Receptor | ^13^ |
| MTDH | -0.08 | Yes | 0.57 | Metadherin | ^14^ |
| FOXM1 | -0.06 | Yes | 0.31 | Forkhead Box Protein M1 | ^15^ |
| SFRP1 | -0.88 | Yes | 0.83 | Secreted Frizzled-Related Protein 1 | ^16^ |
| NES | -0.40 | Yes | 1.75 | Nestin | ^17^ |
| SMAD4 | -0.06 | Yes | 0.88 | SMAD Family Member 4 | ^18^ |
| L1CAM | -0.06 | Yes | 3.38 | L1 Cell Adhesion Molecule | ^19^ |
| CD274 | -0.03 | Yes | 1.39 | Cluster of Differentiation 274 | ^20^ |
| CTHRC1 | -2.29 | Yes | 1.38 | Collagen Triple Helix Repeat Containing 1 | ^21^ |
| ELF5 | 0.00 | Yes | 4.04 | E74 Like ETS Transcription Factor 5 | ^22^ |

**Supplementary Table 3**: Differential expression of iron and lipid metabolism genes involved in ferroptosis identified in mitomiR-3 mimic and sponge transfected TNBC cells.

| **Gene symbol** | **mitomiR-3 Mimic logFC** | **mitomiR-3 Sponge logFC** | **Function in Iron and Lipid metabolism** | **Function in ferroptosis** |
| --- | --- | --- | --- | --- |
| PLCL1 | 1.90 | -7.50 | Membrane Lipid | - |
| STEAP2 | 0.80 | -4.82 | Iron intake | - |
| TF | 0.00 | 2.45 | Iron intake | Inducer |
| CYP4F3 | 0.00 | 2.22 | Fatty Acid | Inducer |
| LIPG | 0.24 | 2.08 | Fatty Acid | Inducer |
| SCD | -0.41 | -1.87 | Fatty Acid | Inhibitor |
| ALOX15 | 0.00 | 1.97 | Unsaturated fatty acids biosynthetic process | Inducer |
| FADS1 | 0.20 | 1.63 | Fatty Acid | Inducer |
| HEPH | 0.00 | -2.13 | Iron efflux | Inhibitor |
| FAR1 | 0.33 | 1.44 | Fatty Acid | Inducer |
| ACSL6 | 0.00 | 2.34 | Fatty Acid | Inducer |
| SLC39A8 | 0.16 | 1.33 | Iron intake | Inducer |
| STEAP4 | -0.08 | 1.31 | Iron intake | Inducer |
| ELOVL7 | 0.29 | 0.83 | Fatty Acid | Inducer |
| FASN | -0.51 | -1.24 | Fatty Acid | Inhibitor |
| CP | 0.87 | -1.03 | Iron efflux | Inhibitor |
| EGLN1 | 0.80 | -0.90 | Iron utilization | Inhibitor |

**Supplementary Table 4**: List of oligonucleotides used.

| **Name** | **Sequence** | **Length** | **Purpose** |
| --- | --- | --- | --- |
| **Linker-S** | TCGAGCTGGTTAACGACGGGTCCCGACGTTTAAACGACG | 39 | Sponge cloning |
| **Linker-AS** | AATTCGTCGTTTAAACGTCGGGACCCGTCGTTAACCAGC | 39 |  |
| **PremiR3-SSp** | GTCCCCCTCTACCTGCCATCTTCCCAAATTCCTCTACCTGCCATCTTCCCAGG | 53 |  |
| **PremiR3-ASSp** | GACCCTGGGAAGATGGCAGGTAGAGGAATTTGGGAAGATGGCAGGTAGAGGGG | 53 |  |
| **pMSCV-For** | TTTATCCAGCCCTCACTCC | 19 | Sponge cloning and PCR |
| **pMSCV-Rev** | TTGTGTAGCGCCAAGTGCC | 19 |  |
| **ZEB1_FP** | CCTGTCCATATTGTGATAGAGGC | 23 | Gene expression |
| **ZEB1_RP** | ACCCAGACTGCGTCACATGT | 20 |  |
| **GPX4_FP** | TACTGCAACAGCTCCGAGTTC | 21 |  |
| **GPX4_RP** | GGTGCCAAAGAAAGAAAGTCC | 21 |  |
| **ACTB_FP** | GACGACATGGAGAAAATCTG | 20 |  |
| **ACTB_RP** | ATGATCTGGGTCATCTTCTC | 20 |  |
| **ZEB1_3’UTR_FP** | ACCTCTCGAGCTTCACAGGTTGCTCCTTCT | 30 | Luciferase cloning |
| **ZEB1_3’UTR_RP** | CACCCCTGCAGGCTGTTAGGCAGTGAGGAATGT | 33 |  |
| **GPX4_2825_For** | CTGCAAGCTCCGCCTAC | 17 | ChIP |
| **GPX4_2826_Rev** | AGACGGGCAGATCACGA | 17 |  |

Supplementary reference

1.	Dongre, A. & Weinberg, R. A. New insights into the mechanisms of epithelial-mesenchymal transition and implications for cancer. *Nat Rev Mol Cell Biol* **20**, 69–84 (2019).

2.	Yang, J. *et al.* Lipocalin 2 promotes breast cancer progression. *Proc Natl Acad Sci U S A* **106**, 3913–3918 (2009).

3.	Qian, X.-L. *et al.* Caveolin-1: a multifaceted driver of breast cancer progression and its application in clinical treatment. *Onco Targets Ther* **12**, 1539–1552 (2019).

4.	Cuevas, E. P. *et al.* LOXL2 drives epithelial-mesenchymal transition via activation of IRE1-XBP1 signalling pathway. *Sci Rep* **7**, 44988 (2017).

5.	Zhang, Y. *et al.* Function of the c-Met receptor tyrosine kinase in carcinogenesis and associated therapeutic opportunities. *Mol Cancer* **17**, 45 (2018).

6.	Chang, J. W. *et al.* Claudin-1 mediates progression by regulating EMT through AMPK/TGF-β signaling in head and neck squamous cell carcinoma. *Transl Res* **247**, 58–78 (2022).

7.	Byles, V. *et al.* SIRT1 induces EMT by cooperating with EMT transcription factors and enhances prostate cancer cell migration and metastasis. *Oncogene* **31**, 4619–4629 (2012).

8.	Luo, Y.-Z., He, P. & Qiu, M.-X. FOSL1 enhances growth and metastasis of human prostate cancer cells through epithelial mesenchymal transition pathway. *Eur Rev Med Pharmacol Sci* **22**, 8609–8615 (2018).

9.	Mannion, A. J. *et al.* Pro- and anti-tumour activities of CD146/MCAM in breast cancer result from its heterogeneous expression and association with epithelial to mesenchymal transition. *Front Cell Dev Biol* **11**, 1129015 (2023).

10.	Schwab, A. *et al.* Zeb1 mediates EMT/plasticity-associated ferroptosis sensitivity in cancer cells by regulating lipogenic enzyme expression and phospholipid composition. *Nat Cell Biol* (2024) doi:10.1038/s41556-024-01464-1.

11.	Zhang, X. *et al.* MACC1 promotes pancreatic cancer metastasis by interacting with the EMT regulator SNAI1. *Cell Death Dis* **13**, 923 (2022).

12.	Price, Z. K. *et al.* Disabled-2 (DAB2): A Key Regulator of Anti- and Pro-Tumorigenic Pathways. *Int J Mol Sci* **24**, 696 (2022).

13.	Lo, H.-W. *et al.* Epidermal growth factor receptor cooperates with signal transducer and activator of transcription 3 to induce epithelial-mesenchymal transition in cancer cells via up-regulation of TWIST gene expression. *Cancer Res* **67**, 9066–9076 (2007).

14.	Wang, Z. *et al.* Metadherin regulates epithelial-mesenchymal transition in carcinoma. *Onco Targets Ther* **9**, 2429–2436 (2016).

15.	Yu, C.-P. *et al.* FoxM1 promotes epithelial-mesenchymal transition of hepatocellular carcinoma by targeting Snai1. *Mol Med Rep* **16**, 5181–5188 (2017).

16.	Qu, Y. *et al.* High levels of secreted frizzled-related protein 1 correlate with poor prognosis and promote tumourigenesis in gastric cancer. *Eur J Cancer* **49**, 3718–3728 (2013).

17.	Hagio, M., Matsuda, Y., Suzuki, T. & Ishiwata, T. Nestin regulates epithelial-mesenchymal transition marker expression in pancreatic ductal adenocarcinoma cell lines. *Mol Clin Oncol* **1**, 83–87 (2013).

18.	Ungefroren, H. *et al.* Elucidation of the Role of SMAD4 in Epithelial-Mesenchymal Plasticity: Does It Help to Better Understand the Consequences of DPC4 Inactivation in the Malignant Progression of Pancreatic Ductal Adenocarcinoma? *Cancers (Basel)* **15**, 581 (2023).

19.	Chen, J., Gao, F. & Liu, N. L1CAM promotes epithelial to mesenchymal transition and formation of cancer initiating cells in human endometrial cancer. *Exp Ther Med* **15**, 2792–2797 (2018).

20.	Brabletz, S., Schuhwerk, H., Brabletz, T. & Stemmler, M. P. Dynamic EMT: a multi-tool for tumor progression. *The EMBO Journal* **40**, e108647 (2021).

21.	Hou, M. *et al.* High expression of CTHRC1 promotes EMT of epithelial ovarian cancer (EOC) and is associated with poor prognosis. *Oncotarget* **6**, 35813–35829 (2015).

22.	Chakrabarti, R. *et al.* Elf5 inhibits the epithelial-mesenchymal transition in mammary gland development and breast cancer metastasis by transcriptionally repressing Snail2. *Nat Cell Biol* **14**, 1212–1222 (2012).
