## Supplemental figures for "Mitochondrial genome-encoded mitomiRs regulate cellular plasticity and susceptibility to ferroptosis in triple-negative breast cancer"

Supplementary Fig. 1

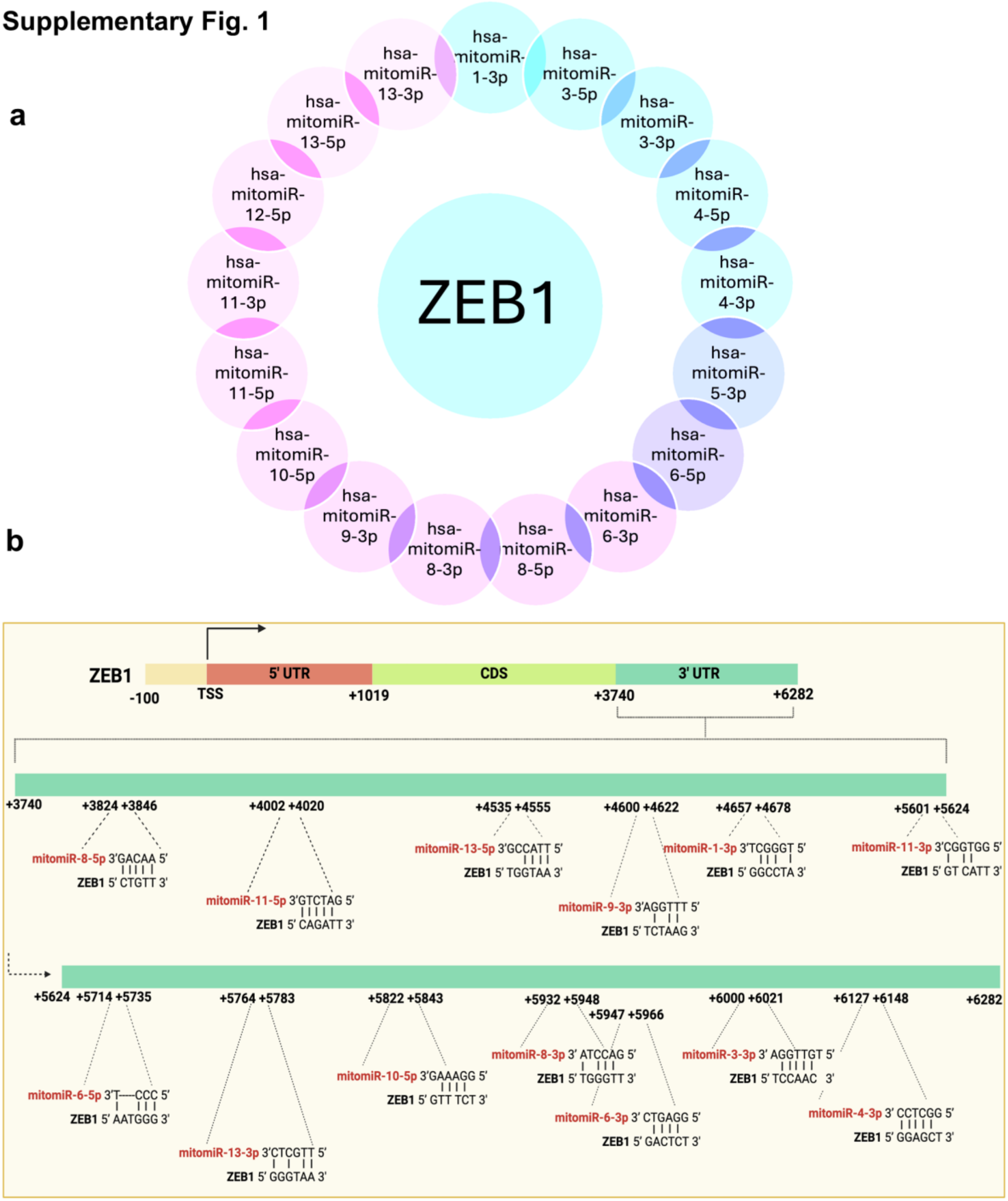


**Supplemental Fig S1**. **Mitochondrial genome encoded miRNAs (mitomiRs) target ZEB1.** (a) Illustration depicting 11 mitomiRs that potentially target ZEB1 predicted using RNA22, RNAhybrid, STarMir and miRanda tools. (b) Schematic representation of predicted binding sites for 11 mitomiRs on the 3'UTR of the ZEB1 mRNA.

**Supplementary Fig. 2**

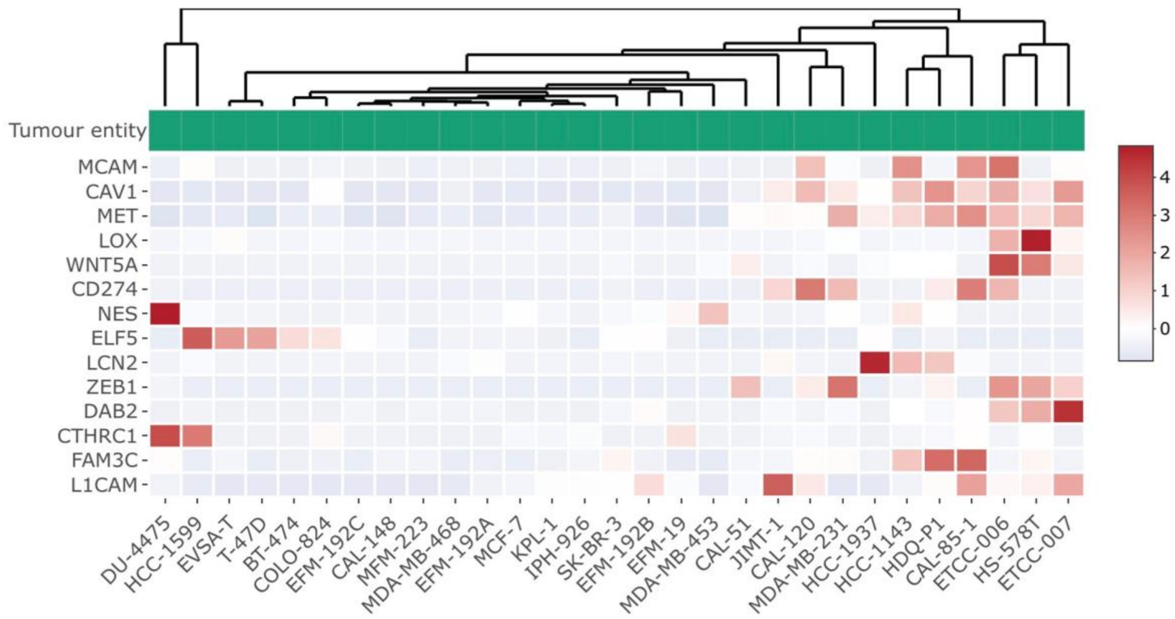


**Supplemental Fig S2. Analysis of** mitomiR-3 target gene expression involved in EMT pathway**.** Heatmap showing expression of EMT pathway genes upregulated in mitomiR-3 sponge cells using public breast cancer cell line RNAseq datasets. TNBC cell lines with mesenchymal features showed high expression of EMT pathway genes upregulated in mitomiR-3 sponge cells.

**Supplementary Fig. 3**

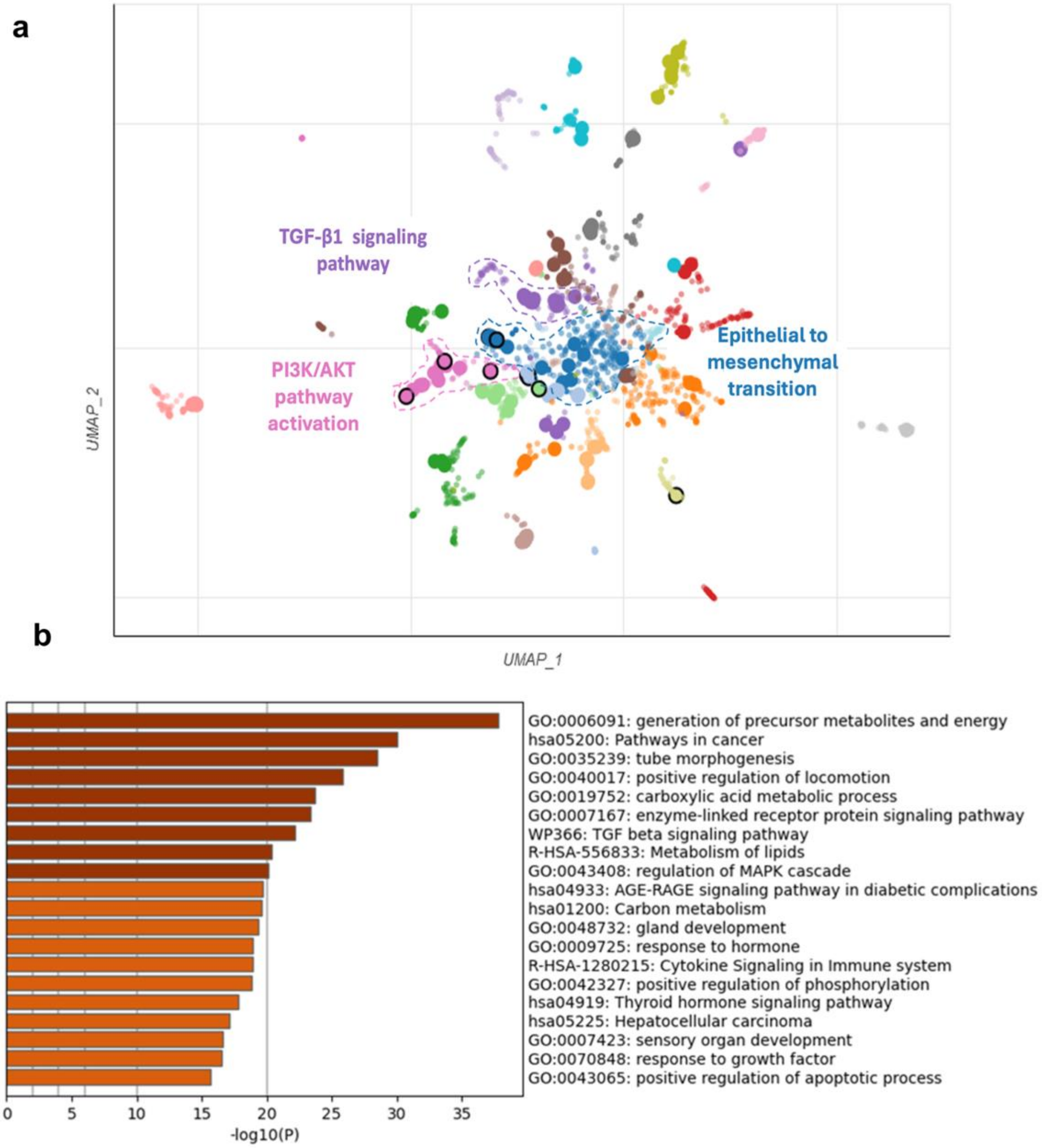


**Supplemental Fig S3. Inhibition of mitomiR-3 induces the expression of mesenchymal marker gene expression.** (a) Upregulated EMT pathway genes from RNA seq-data in mitomiR-3 Sp and Control Sp. Enrichr tool was used to plot the bokeh graph showing the gene enrichment pattern, clustered based on Uniform Manifold Approximation and Projection (UMAP). (b) Bar graph of GO/KEGG enrichment analysis using Metascape.

### Supplementary Fig. 4

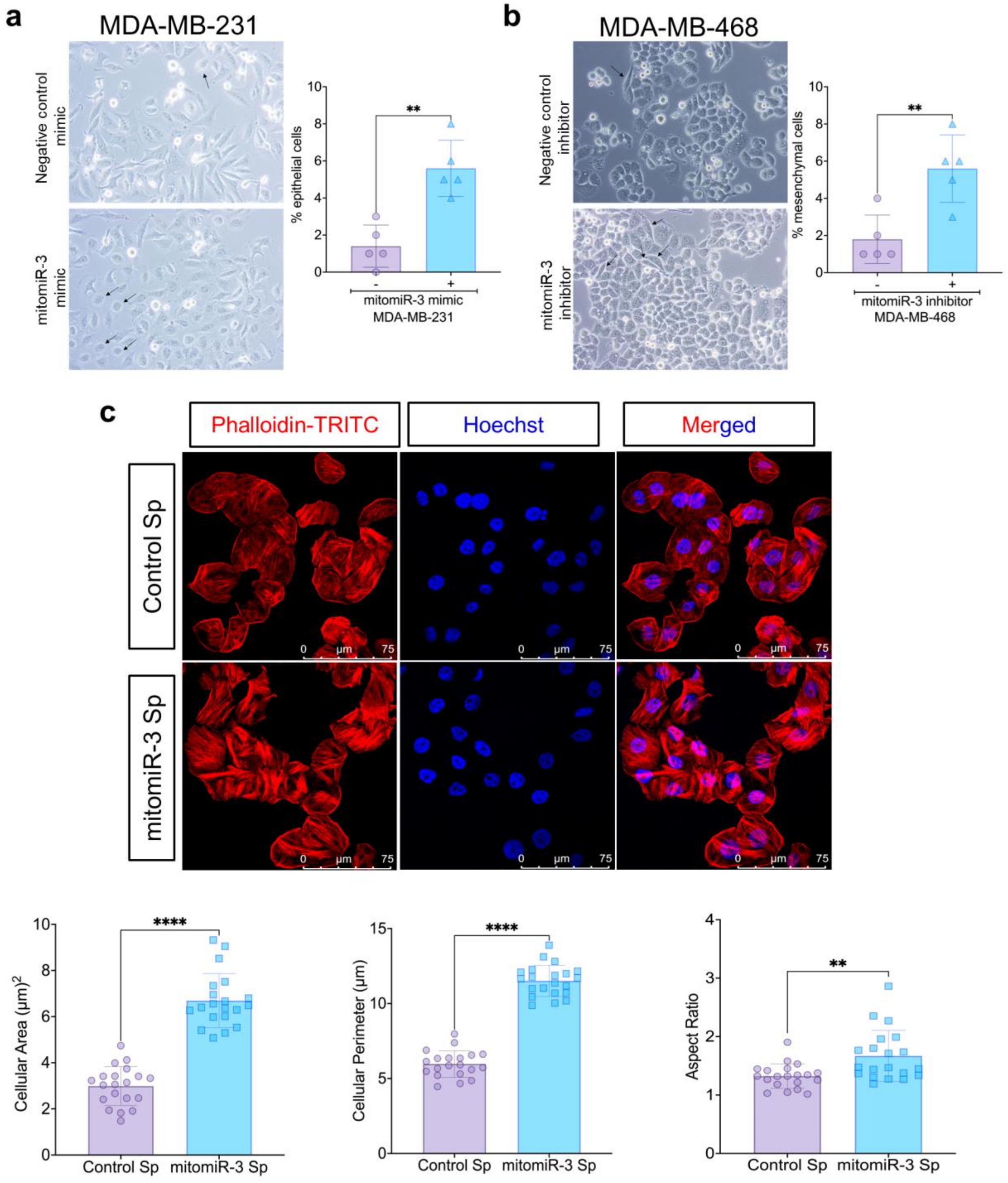


**Supplemental Fig S4. mitomiR-3 loss of function promotes mesenchymal phenotype in mitomiR-3 inhibitor treated and mitomiR-3 sponge TNBC cells.** (a) Bright field microscopy (20x magnification) of the TNBC cell lines MDA-MB-231 transfected with 50nM of mitomiR-3

mimic and its respective negative control oligonucleotides. The bar graph shows the percentage of epithelial cells. MDA-MB-231 (MSL) cells transfected with mitomiR-3 mimic showed an increase in the number of cells showing epithelial phenotype. (b) Bright field microscopy (20x magnification) imaging of the TNBC cell lines MDA-MB-468 transfected with 50nM of mitomiR-3 inhibitor and its respective negative control oligonucleotides. Bar graph showing the percentage of mesenchymal cells. MDA-MB-468 (BL-1) cells transfected with mitomiR-3 inhibitor showed an increase in the number of cells showing mesenchymal phenotype. (c) Representative images of actin phalloidin staining in mitomiR-3 Sp and Control Sp (Scale bars, 50μm). A total of 12 different ROIs were analyzed per group. The bar graph shows an increase in cellular area, cellular perimeter and aspect ratio in mitomiR-3 Sp cells. mitomiR-3 Sp cells showed increased mesenchymal cell type specific features.

**Supplementary Fig. 5**

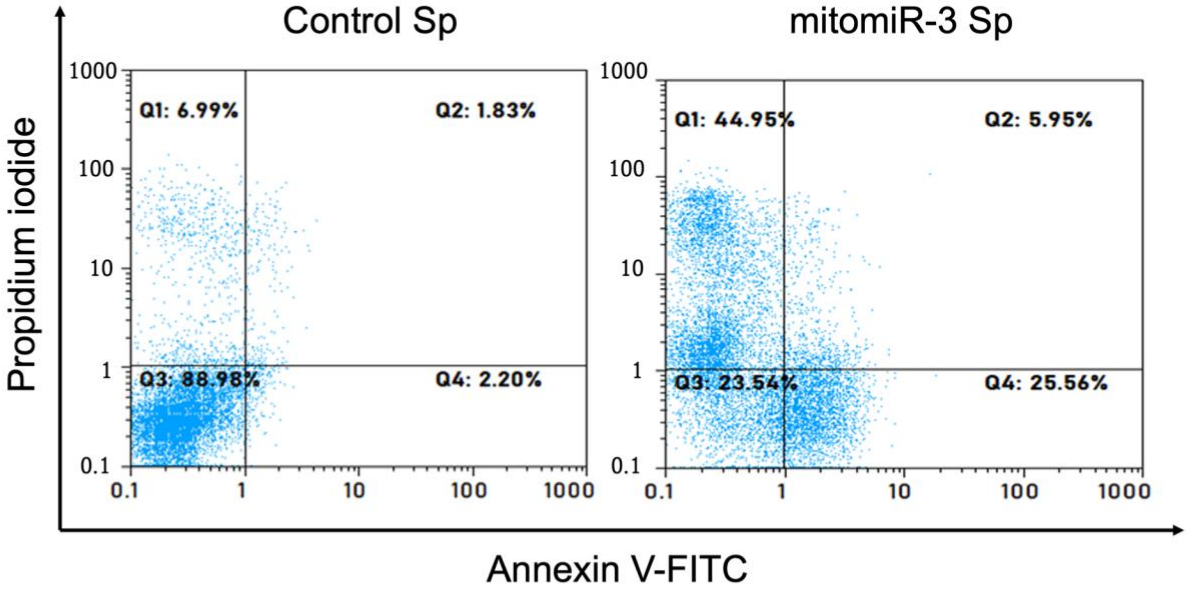


**Supplemental Fig S5. Inhibition of mitomiR-3 increased cell death in mitomiR-3 sponge TNBC cells.** Annexin V-FITC apoptosis assay to assess difference between mitomiR-3 Sp and Control Sp. *p<0.05, **p<0.01, ***p<0.001 and ****p<0.0001.

**Supplementary Fig. 6**

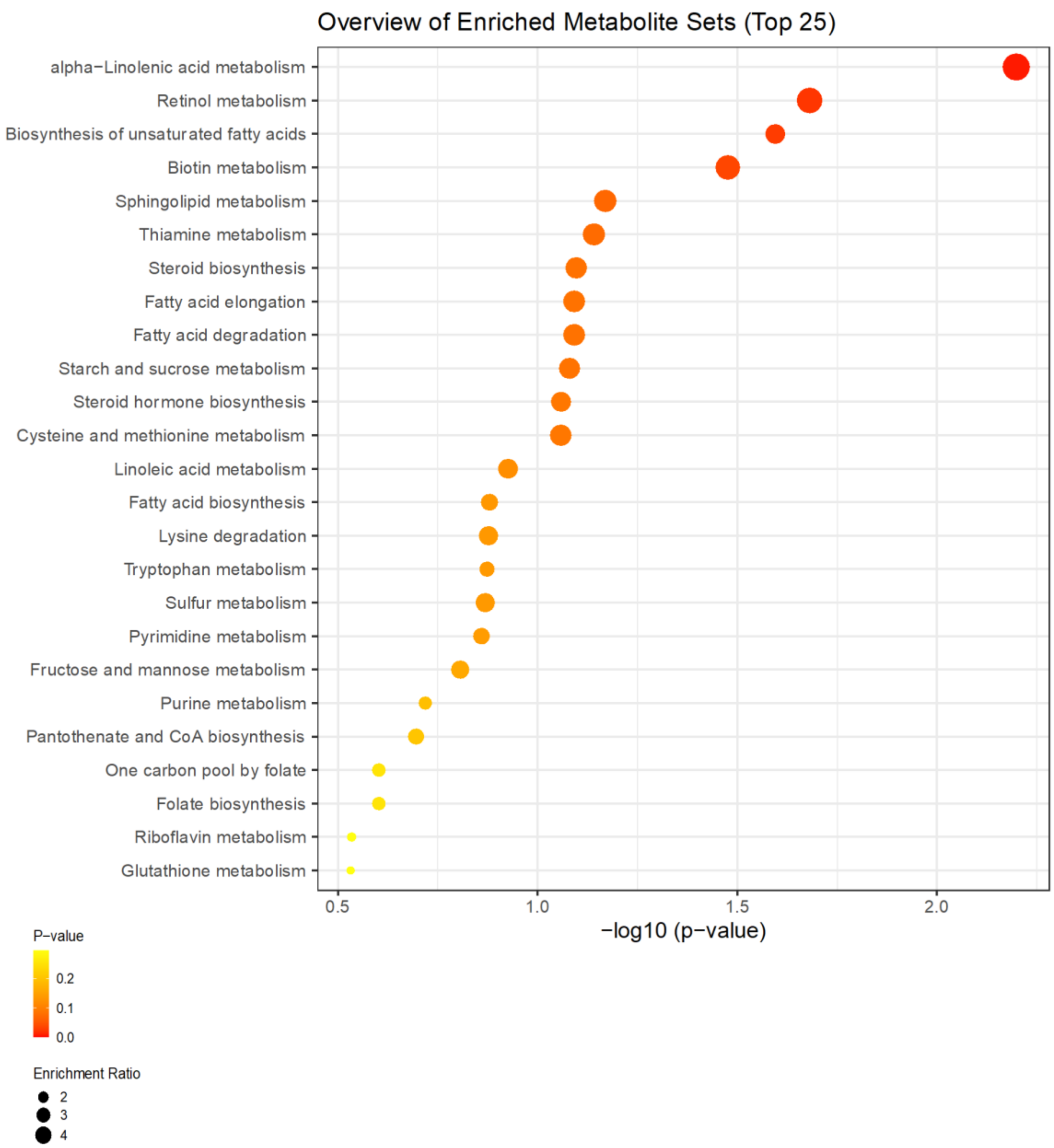


**Supplemental Fig S6. Pathway analysis of metabolomics data showed enrichment of PUFA metabolism in mitomiR-3 sponge TNBC cells.** The bubble plot represents the metabolite set enrichment in mitomiR-3 Sp cells. The plot showed high enrichment for lipid metabolism pathways.

**Supplementary Fig. 7**

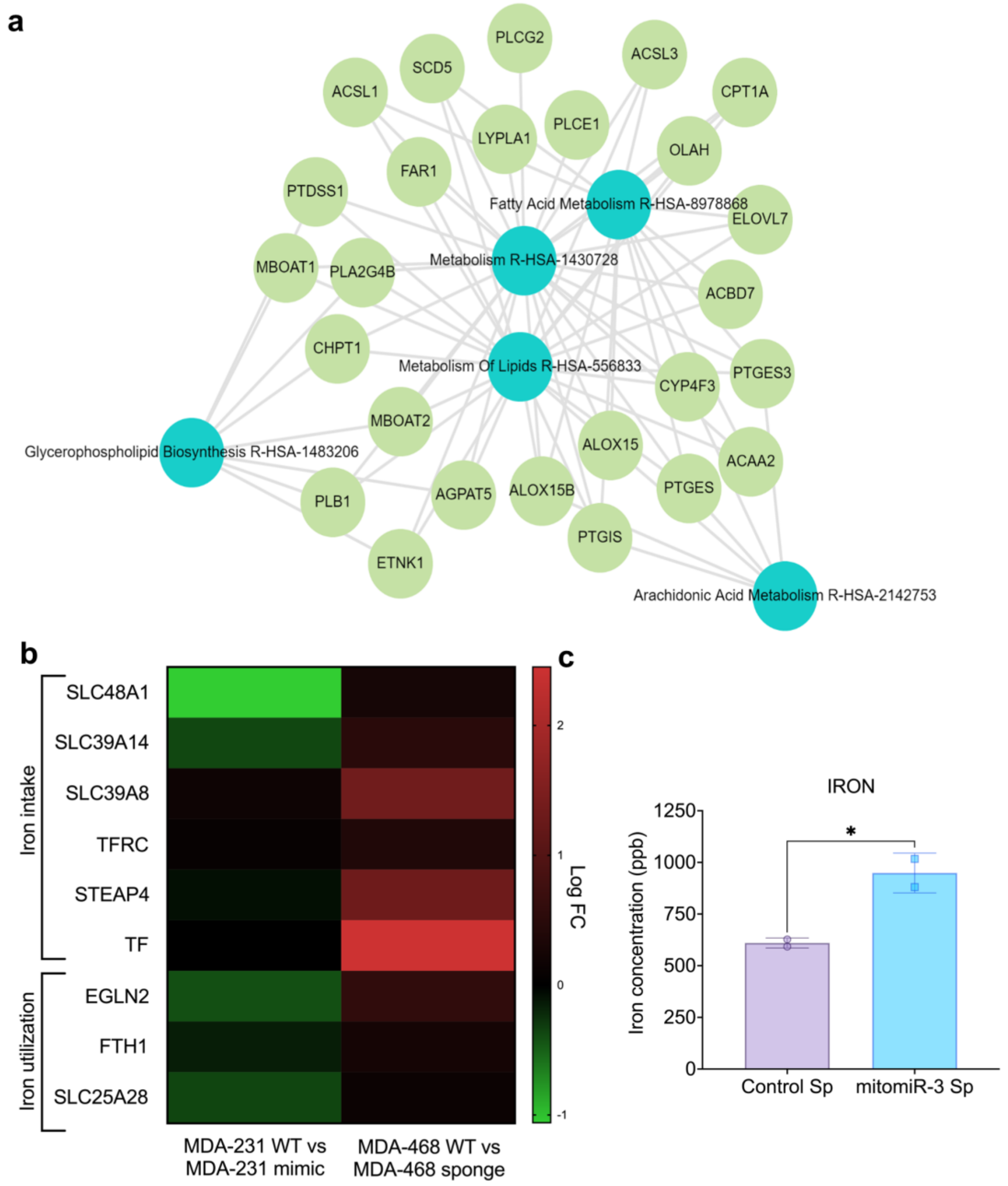


**Supplemental Fig S7. Inhibition of mitomiR-3 reprograms lipid and iron metabolism in mitomiR-3 sponge TNBC cells.** a) Pathway analysis of upregulated genes in mitomiR-3 LOF cells showed enrichment in lipid metabolism. b) Heatmap showing the expression (log FC) of genes associated with iron metabolism in mitomiR-3 Sp and mitomiR-3 mimic. c) Bar plot showing iron content (ppb) in Control Sp and mitomiR-3 Sp. mitomiR-3 Sp had increased accumulation of iron when compared to control Sp. $^{*}p<0.05$.

### Supplementary Fig. 8

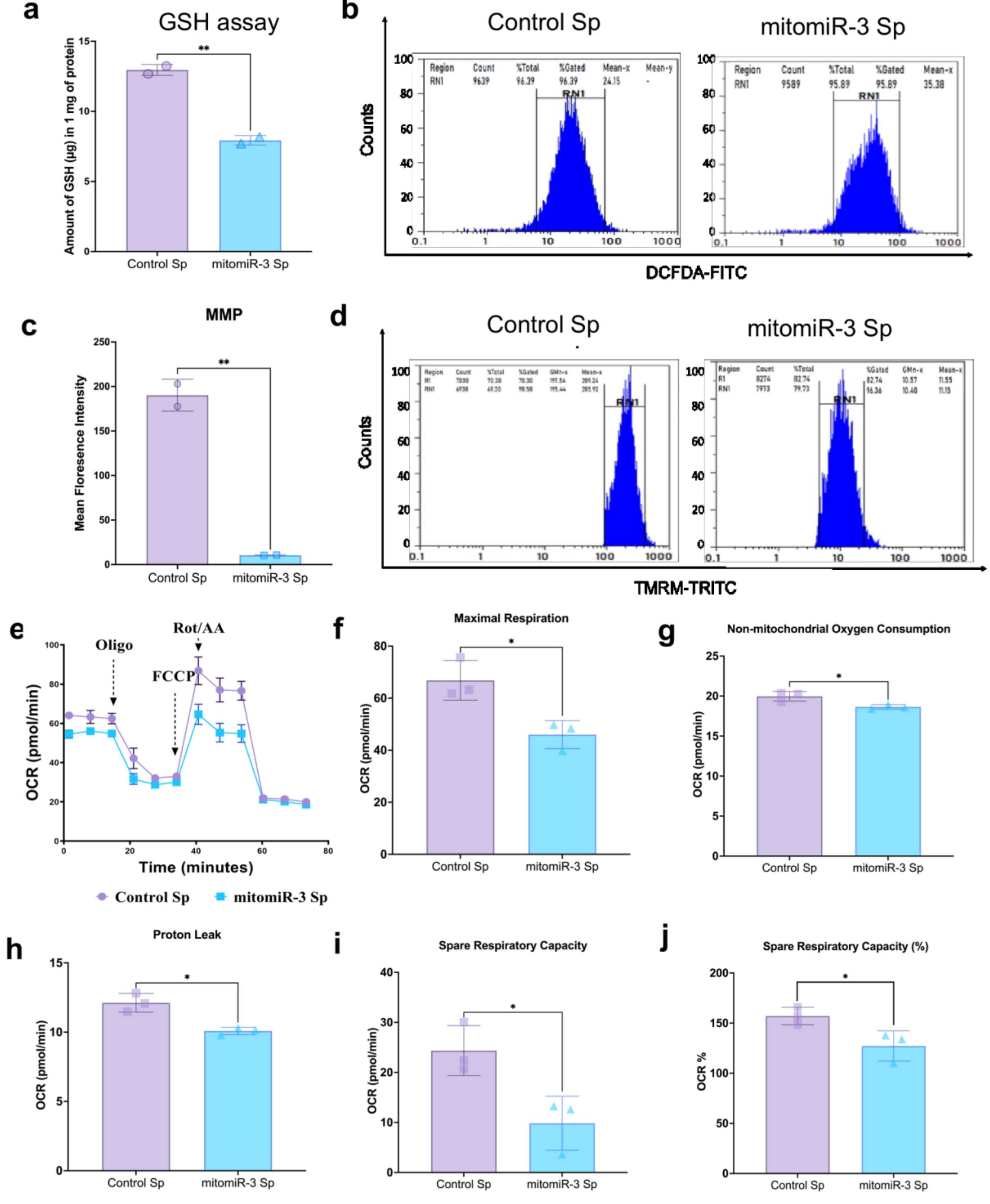


**Supplemental Fig S8. Inhibition of mitomiR-3 reduces cellular glutathione, mitochondrial membrane potential and oxygen consumption rate in mitomiR-3 sponge TNBC cells.** (a) The graph represents intracellular glutathione levels quantified using spectrophotometric glutathione assay in Control Sp and mitomiR-3 Sp. Intracellular GSH is significantly reduced in mitomiR-3 Sp cells compare to the Control Sp cells; (b) Flow cytometry histogram illustrating ROS content by DCFDA staining. (c) Bar graph showing flow cytometric analysis of Control Sp and mitomiR-3 Sp cell lines stained for mitochondrial membrane potential (TMRM). The mitochondrial membrane potential was significantly reduced in mitomiR-3 Sp cells. (d) Flow cytometry histogram illustrating mitochondrial membrane potential by TMRM staining; (e) Representative graph showing the OCR in Control Sp and mitomiR-3 Sp cell lines. Injections of oligomycin, FCCP, and antimycin A and rotenone (A/R) are indicated. (f–j) Representative bar graph quantifying different mitochondrial bioenergetic parameters: maximal respiration, nonmitochondrial oxygen consumption, proton leak, spare respiratory capacity and spare respiratory capacity (%). *$p<0.05$, **$p<0.01$.
